# CaClust: linking genotype to transcriptional heterogeneity of follicular lymphoma using BCR and exomic variants

**DOI:** 10.1101/2024.04.24.590966

**Authors:** Kazimierz Oksza-Orzechowski, Edwin Quinten, Shadi Darvish-Shafighi, Szymon M. Kiełbasa, Hugo W. van Kessel, Ruben A. L. de Groen, Joost S. P. Vermaat, Julieta H. Sepúlveda Yáñez, Marcelo A. Navarrete, Hendrik Veelken, Cornelis A. M. van Bergen, Ewa Szczurek

**Affiliations:** Faculty of Mathematics, Informatics and Mechanics, University of Warsaw, Warsaw, Poland; Department of Hematology, Leiden University Medical Center, Leiden, Netherlands; Cancer Research UK, Cambridge Institute, Cambridge, UK; Department of Biomedical Data Sciences, Leiden University Medical Center, Leiden, Netherlands; Escuela de Medicina, Universidad de Magallanes, Punta Arenas, Chile; Institute of AI for Health, Helmholtz Zentrum München, German Research Center for Environmental Health, Neuherberg, Germany

**Author notes:** These authors have contributed equally to this work (Kazimierz Oksza-Orzechowski, Edwin Quinten, Cornelis A. M. van Bergen, Ewa Szczurek).

**Keywords:** Cancer genetics, tumour heterogeneity, statistical methods, Follicular Lymphoma

## Abstract

Tumor tissues exhibit high genotypic and transcriptional heterogeneity, resulting from tumor evolution and affecting cancer progression and treatment. These two types of heterogeneity in follicular lymphoma were so far predominantly studied in separation. To comprehensively investigate the evolution and genotype to phenotype maps in follicular lymphoma, we introduce CaClust, a probabilistic graphical model that integrates deep whole exome, single-cell RNA and B-cell receptor sequencing data to infer clone genotypes, cell-to-clone mapping, and single-cell genotyping. CaClust outperforms a state-of-the-art model on simulated and patient data. In-depth analysis of 22492 single cells and whole exomes from four follicular lymphoma samples using CaClust gives insights into effects of driver mutations, follicular lymphoma evolution, and possible therapeutic targets. CaClust single-cell genotyping agrees with genotypes observed in an independent targeted resequencing experiment. Our approach is the first to evaluate the strength of genotype to phenotype links in follicular lymphoma in the evolutionary context of the disease.

## Background

From their onset, cancers are subject to continuous evolutionary processes, during which tumour cells acquire mutations in their genomes, forming clones and giving rise to genotypic heterogeneity [1–3]. At the same time, transcriptional heterogeneity is common, with various tumour cell subpopulations having different transcriptional programs. Cancer cell phenotype is thought to be predominantly driven by genetic alterations. However, recent studies suggest that distinct cancer cell states may emerge due to non-genetic factors [4–8]. In this context, a fundamental question arises: to which extent the observed transcriptional heterogeneity in tumours is explainable by the genotypic differences between clones, and which remaining variance should rather be attributed to other effects? A better understanding of the genotype-phenotype link in cancer could guide personalised treatment and prompt development of novel therapies [6, 9–11].

Follicular lymphoma (FL) is a malignancy of mature B-cells that are arrested at the germinal center stage and usually presents as pathological lymph nodes [12, 13]. FL is a paradigmatic disease with a clinical course that can range from spontaneous regression or stable disease for years, to transformation into aggressive diffuse large B cell lymphoma. In addition to acquisition of mutations in BCR loci, aberrant somatic hypermutation in non-BCR loci causes accumulation of somatic variants that can be clonal (early events, present in all clones) or subclonal (later events, specific to a subset of clones) [14]. Under the assumption that acquisition of a potential oncogenic subclonal mutation occurs alongside BCR diversification, BCRs can serve as markers in the study of clonal evolution in FL tumours [15]. At the same time, FL cells may display transcriptional heterogeneity, with different transcriptional subpopulations displaying varying drug responses [16]. An example of a genotype to transcriptional phenotype link in FL is the presence of N-linked glycosylation motifs in BCR, which drives FL cells from a more light zone-like gene expression profile toward a dark zone-associated transcriptional program [17]. Still, the genotype to phenotype maps of FL have not been so far comprehensively studied.

The major obstacle in investigating the relation between genotypic and transcriptional heterogeneity in tumours is the fact that simultaneous DNA and RNA profiling of single cells is not possible using established experimental protocols. Indeed, innovations in this area emerged only very recently and are not commercially available [18–20]. To address this, computational methods were proposed that probabilistically match genomic alterations between bulk DNA sequencing and single cell RNA sequencing (scRNA-seq) or spatial transcriptomics data [15, 21– 25]. In particular, our approach, CACTUS was previously applied to FL data by clustering cells by BCR sequences and performing cluster-to-clone assignment by mutation matching, benefiting from BCR information in this task [15]. However, the clustering of cells in CACTUS was effectively limited to grouping of cells with identical BCRs and required defining a hyperparameter corresponding to the (unknown) number of clusters. As an alternative to computational approaches, targeted DNA resequencing of single cells previously sequenced using scRNA-seq can also be used for genotyping cells [26]. Unfortunately, due to technical limitations it can only be performed with a very small number of variants, and is subject to noise in the scRNA data due to bursty gene expression.

Here, we use deep whole exome sequencing (WES), single cell RNA sequencing, single cell BCR sequencing (BCR-seq) and probabilistic modelling to deeply investigate the nature of tumour evolution as well as genotype to transcriptional phenotype interactions in four FL samples. To this end, we propose CaClust, a nonparametric Bayesian extension of CACTUS [15], which is able to find a confident cell-to-clone mapping, improve estimation of clone genotypes, and infer genotypes for single cells. CaClust identifies the number of BCR clusters from the data and models probabilistic profiles of BCR sequences characteristic of every cluster. Applying CaClust to FL samples we find a range of genotype to phenotype links of varying strength. Focusing on the sample with the largest strength, we investigate mutations that could trigger specific transcriptional phenotypes. Additionally, we uncover the evolutionary link between two time-separated samples from another FL patient. In summary, our analysis enables comprehensive analysis of FL cell phenotypes in the context of their clonal origin, uncovering underlying tumour evolutionary mechanisms and targetable dependencies.

## Results

### Approach overview

We performed comprehensive molecular profiling of three FL patients: K4B and K5B, two samples from 69 year old male subject S8934 separated by three years, sample K6B from 32 year old male subject S13530, as well as K7B from 48 year old woman S11770 (Fig. 1 1a). Each sample was profiled using WES at 1500x coverage. scRNA-seq and heavy and light chain scBCR-seq resulted in a total of 22492 single cells sequenced at average 1620 transcriptome-wide genes, and average 24 full length umis of the expressed BCR genes per single cell.

**Figure 1:**
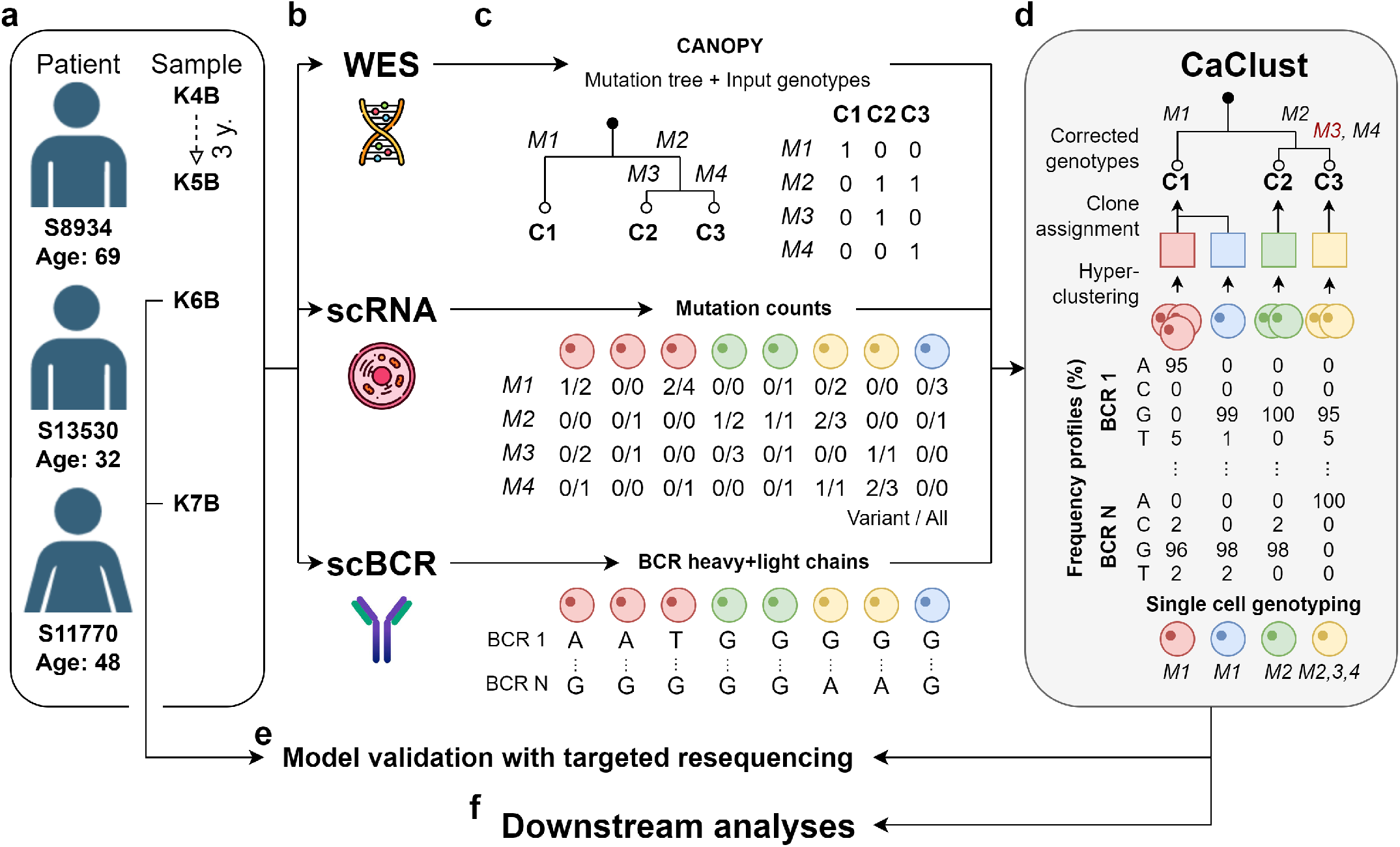
Application of CaClust in this study: **a** 4 samples from 3 FL patients were chosen for inclusion in the study; **b-c** data collection and preprocessing; **d** model application to infer the clone genotypes, BCR hyperclustering with assignment, and clone clusters, output cell genotypes are obtained with the mapping of cells to their clone of origin; **e** after model application an additional resequencing experiment was performed on samples K6B and K7B to validate the output cell genotypes; **f** the output cell genotypes and clone structure were used in downstream analyses.

In the CaClust model application pipeline, we first perform variant calling and copy number (CN) analysis on WES data, and next we use their output to estimate the input phylogenetic tree of the clones and their genotypes. From the scRNA-seq data, after standard alignment and mapping, we extract for each cell the variant and total read counts at the single nucleotide variant (SNV) positions. From the BCR-seq data we extract the BCR sequence for each cell, comprising of the concatenated heavy and light BCR chains (Fig. 1c).

From this input, CaClust aims to simultaneously reconstruct *BCR hyperclusters* of cells while assigning those hyperclusters to tumour clones (Fig. 1d). Each BCR hypercluster is represented by its frequency profile of BCR nucleotides at different positions. We assume all cells in a hypercluster must come from the same clone. The tumour clones are represented by the SNVs present in the clone’s genotype. The BCR hyperclusters are assigned to the clones by a probabilistic matching of variant reads at the SNV positions. Finally, *clone clusters* of cells are identified by tracking the clone that the BCR hypercluster of each cell is assigned to. Effectively, the cells are mapped to clones based on their shared BCR frequency profile characteristic of their hypercluster, as well as the genotype of its assigned clone, which then we also use to perform single cell genotyping (Fig. 1d). In this way, CaClust marries genotypes with phenotypes and enables detailed gene expression analysis of clones.

The output cellular genotypes obtained with our genotype-to-phenotype mapping were validated using simulated data and an independent resequencing experiment (Fig. 1e) and next used for downstream analyses of the evolutionary structure and the genotype to phenotype links in the FL samples (Fig. 1f).

### CaClust model validation

Before performing detailed downstream analyses, we validated the performance of the CaClust model on simulation scenarios with known ground truth and evaluated its quality on the FL samples, comparing against a predecessor model, as well as checked the correctness of its single cell genotyping with an independent targeted resequencing experiment.

We devised eight simulation scenarios (see Methods) varying three properties of a FL dataset: the scRNA read depth, the number of BCR hyperclusters, and the variance of BCR sequences within a hypercluster. A scenario with parameter values resembling the FL patient datasets used in this study was established as a baseline (referred to as *Basic*) and by varying one of these properties eight further scenarios were created with: i) high and ii) low number of scRNA reads; referred to as *High reads* and *Low reads*, respectively; iii) sparse BCR clustering (resulting in more BCR hyperclusters, referred to as *Sparse clusters*); iv) high variance BCR sequences within hyperclusters (*High variance BCR*); and finally, four scenarios with centroid behaviour, where a portion *x* of cells within a fraction *y* of hyperclusters BCR sequence identical to the most probable of its hypercluster’s BCR profile (in four versions of (*x, y*) v-viii): (0.8, 0.8), (0.8, 0.2), (0.2, 0.8), (0.2, 0.2); referred to as *x centroids in y clusters*). The simulation process and all scenario details are described in Methods. Both CaClust and its predecessor model CACTUS were applied to ten datasets simulated per each scenario and their performance was evaluated using three metrics: cell to clone assignment accuracy, accuracy of clone genotype reconstruction, and the quality of the hyperclustering reconstruction (see Methods).

CaClust outperformed CACTUS in all scenarios and showed near perfect scores in the Basic type, High reads, as well as centroid scenarios (Fig. 2a-c). In terms of cell to clone assignment accuracy, the only scenarios that affected its performance were Low reads, sparse clusters and High variance BCRs. However, in all these scenarios CaClust significantly improved over CACTUS and kept median accuracy over 0.64 (Fig. 2a). In the task of genotype reconstruction CaClust showed a less pronounced improvement over CACTUS (Fig. 2b). Both models performed near perfect in reconstructing the true clone genotypes, with better reconstruction accuracy in scenarios with high scRNA read counts and worse accuracy in scenarios with degraded (sparse or high variance) BCR clustering. Hyperclustering reconstruction comparison again demonstrated a major advantage of CaClust (Fig. 2c). It achieved high adjusted rand index (ARI) scores in all but the high variance BCR scenario, which is specifically designed to give almost no BCR information. CACTUS performed poorly in all scenarios with basic and degraded BCR structure, since it did not manage to reconstruct hyperclusters with varied BCR sequences. However, in the scenarios with centroid behaviour, which can happen for real BCR data, its performance improves, albeit not to the level of CaClust.

**Figure 2:**
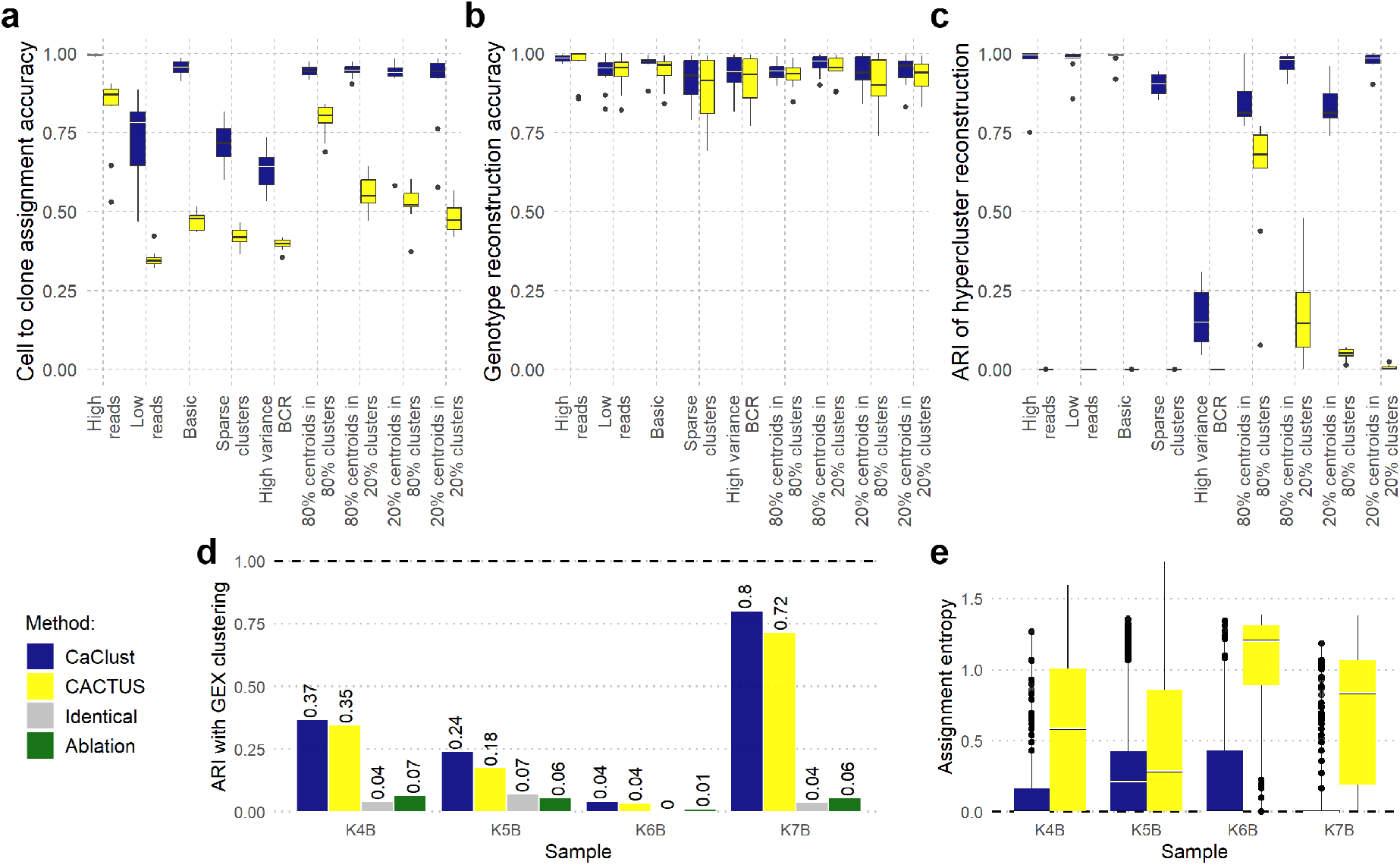
CaClust validation results: **a-c** performance comparison on simulation scenarios vs. predecessor model CACTUS; **d-e** comparison on experimental data: **d** ARI agreement with gene expression clustering for CaClust clones, CACTUS clones, clusters of cells with identical BCR sequences, and hyperclusters from an ablation study that does not use scRNA variant data; **e** the assignment entropy of cells to clones in CaClust and CACTUS results. Bold dashed line: the value for the best possible clustering agreement as measured with ARI (**d**) and the value of the least entropy corresponding with the highest model certainty of assignment (**e**).

Next, we compared performance of CaClust and CACTUS methods on the FL patient sample datasets by their sensitivity to the possible link between clone genotypes and transcriptional heterogeneity, measured using ARI against an independent cell clustering by gene expression (see Methods and Supplementary Figure 1). We used two naive approaches as baselines: grouping cells with identical BCR sequences; and an ablation study with a stripped-down version of the CaClust model, which only produces hyperclusters with no further grouping into clones. CaClust achieved higher agreement with gene expression clustering as compared to CACTUS for all FL samples (Fig. 2d). This indicates that the clustering of cells into clones found using CaClust identifies clones that are more distinct on the phenotypic level. Both models obtained higher ARI than the baselines, showing that the integration of both BCR and scRNA variant data is more accurate in finding phenotypically distinct populations of cells.

Further, we compared CaClust and CACTUS by their confidence of cell to clone assignment, measured with entropy. CaClust assigned cells to clones with lower entropy than CACTUS, indicating that the model was better able to extract information from the integrated data and reduce noise to make confident predictions (Fig. 2e, Supplementary Figure 2).

Finally, we performed an additional targeted resequencing of samples K6B and K7B for independent experimental validation of cell genotyping performed by CaClust (see Methods). CaClust single cell genotypes show *>* 90% agreement for 4*/*8 resequenced variants, which increases to 6*/*8 variants when accounting for random monoallelic expression in the resequencing data (see Supplementary Notes and Supp. Tabs. 2, 3).

### FL samples show different strengths of genotype-phenotype influence

To inspect the level of genotypic and transcriptional heterogeneity, and to assess the strength of genotype to phenotype link in the four analysed FL samples, we investigated the number and proportion of inferred clones, BCR hyperclusters, as well as visual agreement between the gene expression similarity and clone assignment (Fig. 3).

**Figure 3:**
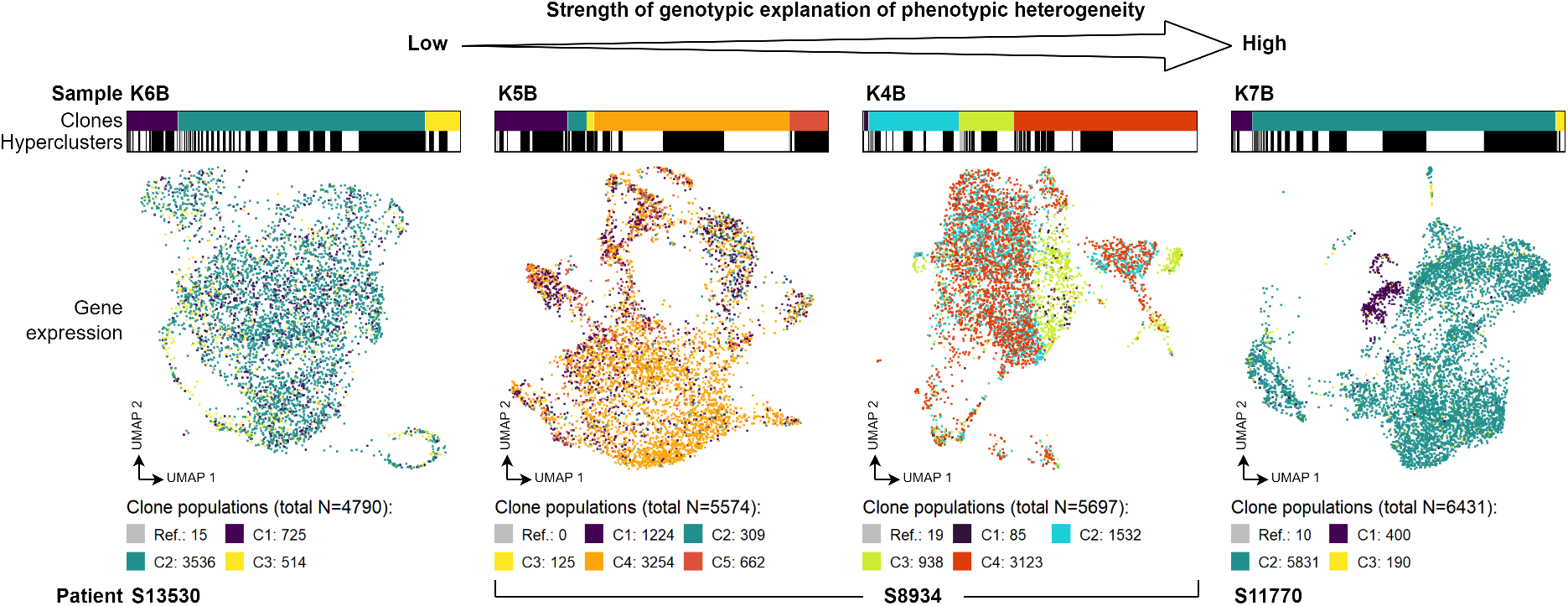
Summary of observed strength of genotype-phenotype link in studied samples. Bars above UMAP plots show clone assignment of cells (colour coded) grouped by their hypercluster assignment (each solid white or black bar is one hypercluster). Cells in UMAP plots of gene expression are also colour coded by clone assignment. Clone populations are provided in the legend. The clones are not shared between samples. The samples are ordered by apparent influence of the clone genotypes on their phenotypic heterogeneity.

Sample K6B from patient S13530 displayed relatively uniform gene expression across its cells. The clustering of cells into the three clones with confidently assigned mutation profiles (see Supplementary Data 1) and 97 identified BCR hyperclusters did not coincide with gene expression similarity (Fig. 3 left). This is in agreement with the lowest *ARI* = 0.04 obtained for that sample (compare Fig. 2d).

In time-separated samples K4B (with four clones and 82 BCR hyperclusters) and K5B (five clones and 32 BCR hyperclusters) coming from patient S8934 we observed higher phenotypic variance, with K4B showing more agreement between the transcriptional subpopulations and the inferred clones (*ARI* = 0.37 vs. *ARI* = 0.24, Fig. 2d; Fig. 3 middle). This could be the effect of specialisation in K5B, where its cells are more genetically homogeneous after genetic selection from K4B.

Sample K7B from patient S11770 (three clones and 92 BCR hyperclusters) displayed the strongest genotype-phenotype link (*ARI* = 0.8, Fig. 2d; Fig. 3 right), with clone C1 forming a separate expression cluster from clones C2 and C3. This points at subclonal variants highly affecting their carriers expression profiles.

In summary, our analysis revealed the clonal and BCR hypercluster structure of the FL tumour samples, and allowed ranking them by the strength of genotypic explanation of transcriptional phenotypic heterogeneity.

### In depth investigation identifies four potential mutations driving clone phenotypes in patient sample K7B

To showcase the usefulness of our approach in the study of the genotype-phenotype link, we performed a detailed analysis of the results in patient sample K7B.

CaClust identified three tumour clones, with phenotypes showing the effects of known mutations. The output clone genotypes (Fig. 4**a**) included four subclonal mutations for which we predicted a pathogenic effect (see Methods): (i) *MYD88* (L265P) mutation, (ii) a variant in the *TSPAN33* gene (both shared between clones C2 and C3), (iii) a missense variant in the HAT domain of the *CREBBP* gene (specific to clone C2), and (iv) an early stop variant *VMA21* (R93X) (specific to clone C1). Clones C2 and C3 shared a larger number of SNVs and appeared to be evolutionally closer to each other than to clone C1.

**Figure 4:**
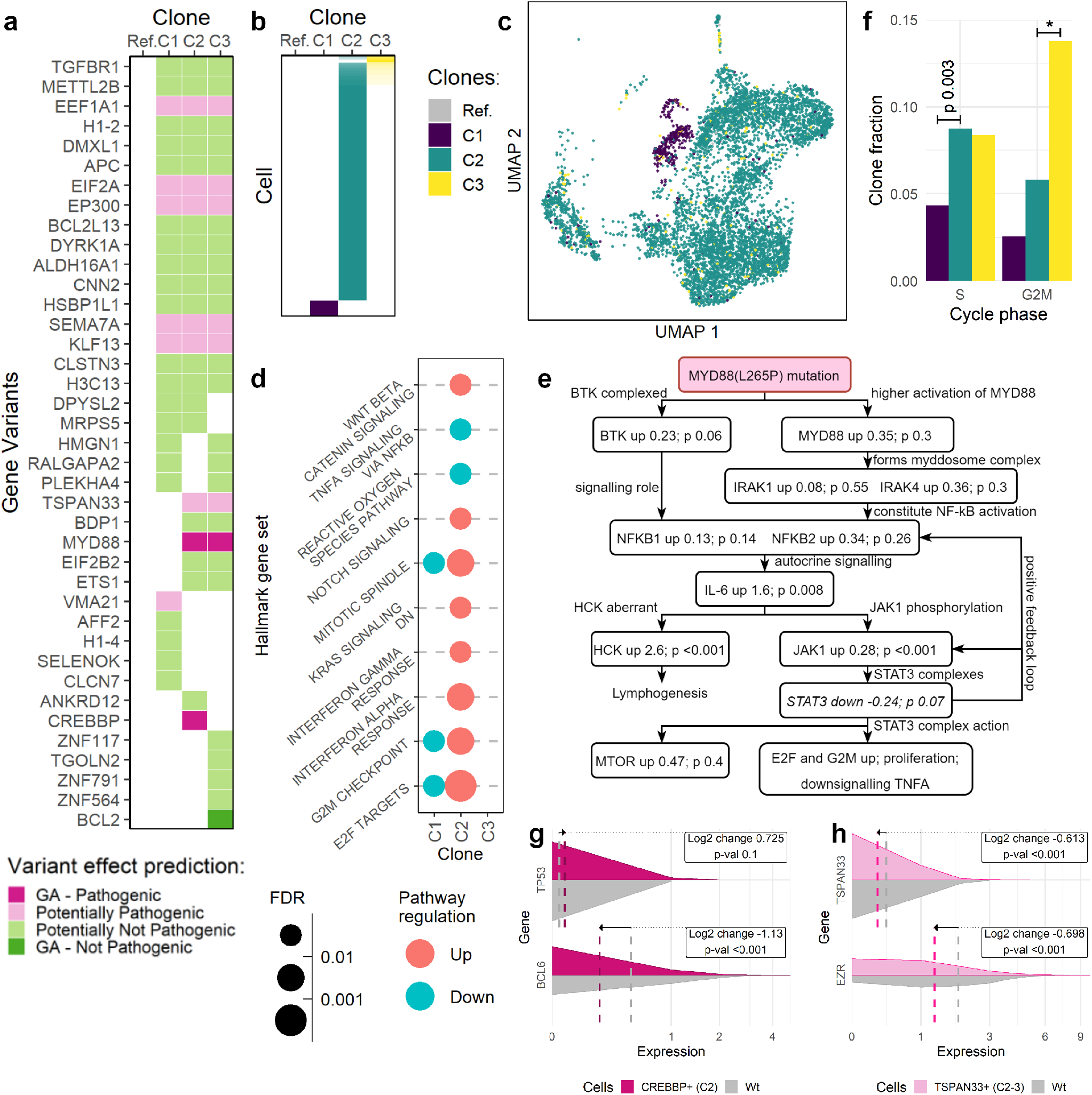
Analysis of CaClust results on sample K7B: **a** output clone genotypes; **b** output cell-to-clone assignment probabilities, higher probability denoted with higher opacity; **c** UMAP reduction of cells’ gene expression, coloured by clone assignment; **d** results of GSEA analysis; **e** graph of predicted *MYD88* (L265P) effects compared with observed DE results; **f** distribution of cells in clones across cell cycle phases; **g** estimated expression of *TP53* and *BCL6* genes in CREBBP+ cells vs. wildtype cells; **h** estimated expression of *TSPAN33* and *EZR* genes in *TSPAN33*+ cells vs. wildtype cells. *–*p <* 0.001, Wt–wildtype.

The model assigned the cells to clones with a very high confidence (Fig.4**b**). Only a small fraction of cells had their assignment probability mixed between C2 and C3. This was in line with the aforementioned higher genotypic similarity of C2 and C3. Among the resulting clone clusters, clone C2 was the most prevalent, with C1 and C3 being smaller in size.

When the UMAP reduction of cells’ gene expression was overlaid with clone clusters (Fig. 4**c**; reproduced from Fig. 3), clone C1 formed its own clear expression cluster, whereas clones C2 and C3 appeared more mixed. This hints at gene expression being highly affected by a subclonal variant, either one that is characteristic to C1 or shared between C2 and C3.

To study the effects of subclonal variants, we first performed differential gene expression analysis between each clone and the rest (see Methods), finding 918 differentially expressed genes. Next, to check for enriched pathways we performed gene set enrichment analysis (Fig. 4**d**). Clone C2 showed upregulated pathways linked to cell proliferation (E2F targets, G2M checkpoint, mitotic spindle (respective FDRs: *<* 0.001, 0.004, 0.007; all FDRs estimated with gene label reshuffling in GSEA) and downregulated tumour necrosis factor alpha signalling (FDR 0.09), both hinting at gained advantages over the other clones.

Next, we investigated whether the known effects of the *MYD88* (L265P) mutation could be observed. Based on a comprehensive study [27], we created the graph of the predicted effects of that mutation in tumour cells as compared to healthy B-cells, and indicated their agreement with our own data (Fig. 4**e**). Most observed effects agreed with the known ones, with the exception of the downregulated *STAT3* expression (logfoldchange −0.236, adj. p-val 0.07, all log fold-change p-values obtained with LRT from DESeq2, see Methods). However, JAK1 affects the formation of the STAT3 complex through protein-protein interaction, and the *STAT3* downregulation in expression may be independent of that process. Moreover, the known effects relate to the comparison of healthy B-cells with *MYD88* (L265P) carriers, whereas our analysis compares tumour cells with multiple additional mutations against each other, which can introduce confounding effects. Since the ultimate effect of the *MYD88* (L265P) is increased tumour proliferation and cell survival, we also analysed the cell cycle distribution of clones (Fig. 4**f**). As expected, cells in C2 and C3 that harbour this mutation proliferated faster, as measured by the fraction of the cells in those clones entering the S phase (hypergeometric test: *p* = 0.003). Additionally, since increased proliferation should require higher energy production, we analysed the enrichment of the beta-oxidation pathway in clones C2 and C3, and found 20*/*25 genes to be upregulated (see Supplementary Figure 3).

We next investigated whether cells assigned to clone C2 follow the behaviour known for carriers of the CREBBP variant. CREBBP proteins with a missense variant in the HAT domain are unable to acetylate the tumour suppressor TP53 and BCL6 oncogene, preventing the tumour suppression mechanisms [28]. As expected, cells assigned to clone C2, even though they express more *TP53* than cells in clones C1 and C3 (log fold-change: 0.725, *p*_*adj*_ = 0.1) and less *BCL6* (−1.13, *p*_*adj*_ *<* 0.001; Fig 4**g**), which should lead to cell cycle arrest, pass through the G2M check-point normally. In contrast, cells from clone 3 are stuck in in the G2M phase (hypergeometric test: *p <* 0.001; Fig. 4**f**).

The TSPAN33 protein has been linked to a migratory phenotype in the B-lymphocytes. It forms complexes on the B-cell membrane with the EZR protein, and its overexpression was linked to an increase in B-cell migration by Navarro-Hernandez *et al*. [29]. In clones C2 and C3 showing the *TSPAN33* missense variant, we observed a decrease in the expression of both *TSPAN33* and *EZR* (log fold-changes: −0.613, −0.698, *p*_*adj*_ *<* 0.001, Fig. 4**h**), which could point to a decrease in migratory capabilities over C1.

The observed *VMA21* (R93X) mutation and was shown by Wang *et al*. [30] to result in a targetable survival dependency. Specifically, VMA21 is a chaperone protein that takes part in the V-ATPase assembly and the mutation p.R93X results in a premature stop and a loss of the C terminus at AA93-101. Consequently, VMA21 is mislocated to the lysosomes, leading to impaired V-ATPase ability to acidify lysosomes that is compensated by an increase in autophagic flux. In the Wang *et al*. study, treatment of *VMA21* (R93X) B-cells with an inhibitor of autophagy regulating ULK1 kinase complex led to their death, while wildtype B-cells remained mostly unaffected; thus, the clone C1 exhibiting *VMA21* (R93X) could also be targetable by such a therapy.

In summary, our analysis firstly, identified which effects of known subclonal mutations can be observed in the phenotypes of specific clones; secondly, pointed to the effects of potentially pathogenic subclonal mutations, which have not been studied yet; lastly, can potentially guide the treatment choice by identifying clones with mutations known to be susceptible to targetable therapy.

### Inferred clone genotypes suggest evolutionary history of the time-related samples

To further showcase the usefulness of our method in studying tumour evolution, we investigated the relation between samples K4B and K5B. Both of those samples were taken from patient S8934, 3 years apart. In CaClust results for sample K4B, we found evidence for a possible founder clone of the tumour in sample K5B, with both genetic and phenotypic similarity.

Firstly, we investigated the 69 SNV variants that were common for both samples and were included in the model analysis (Fig. 5**a**). Variant probabilities for each clone of K4B indicated that clone C3 contained almost all of the common variants with a high probability, while the other K4B clones were missing many of them. Most of those variants in the K5B sample were clonal (i.e., present in all clones), which is expected for a later, possibly descendant tumour sample. This result suggests the possible descendance of all clones found in the later K5B sample from clone C3 or its close ancestor in the evolution of the earlier K4B sample. However, three variants (two in *H1-4* gene and one in *H1-2*) were absent from clones C3 and C4. Those SNVs belonged to a region (chr6 positions 26056K–26156K) that, based on the WES CNA analysis of sample K5B, was found to be affected by a copy number alteration (deletion), which could have happened in the course of evolution of clones C3 and C4 in K5B.

**Figure 5:**
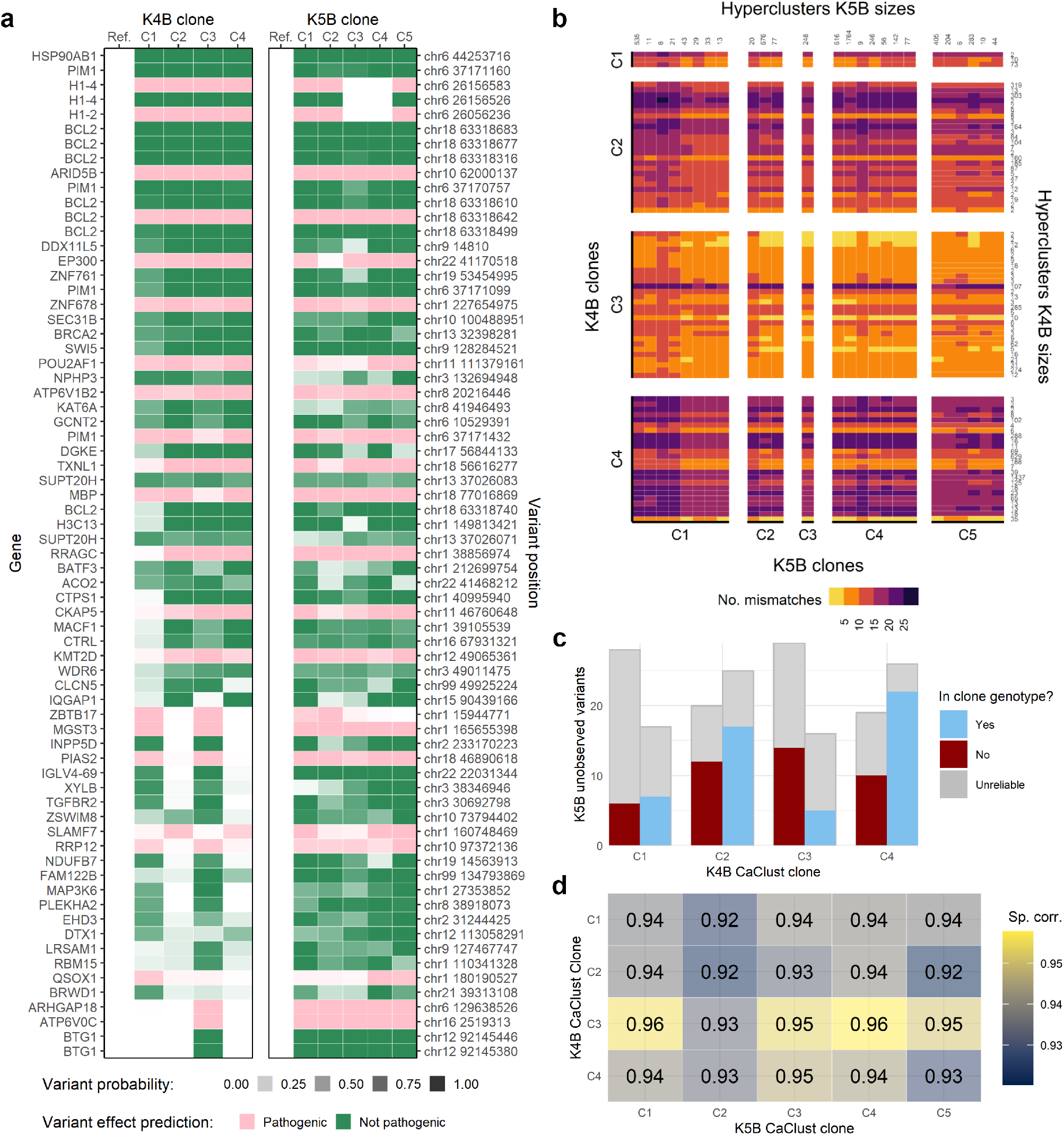
Analysis of the link between K4B and K5B samples using CaClust results: **a** output probabilities of variants common between the samples to be present in clone genotypes; **b** Hamming distance between BCR hyperclusters in K4B and K5B, sorted by their clone assignments; **c** K4B variants unobserved in K5B, by their presence or absence in K4B clone genotypes; **d** Spearmann correlation between gene expression of K4B and K5B clones. Sp. corr.–Spearmann correlation.

**Figure 6:**
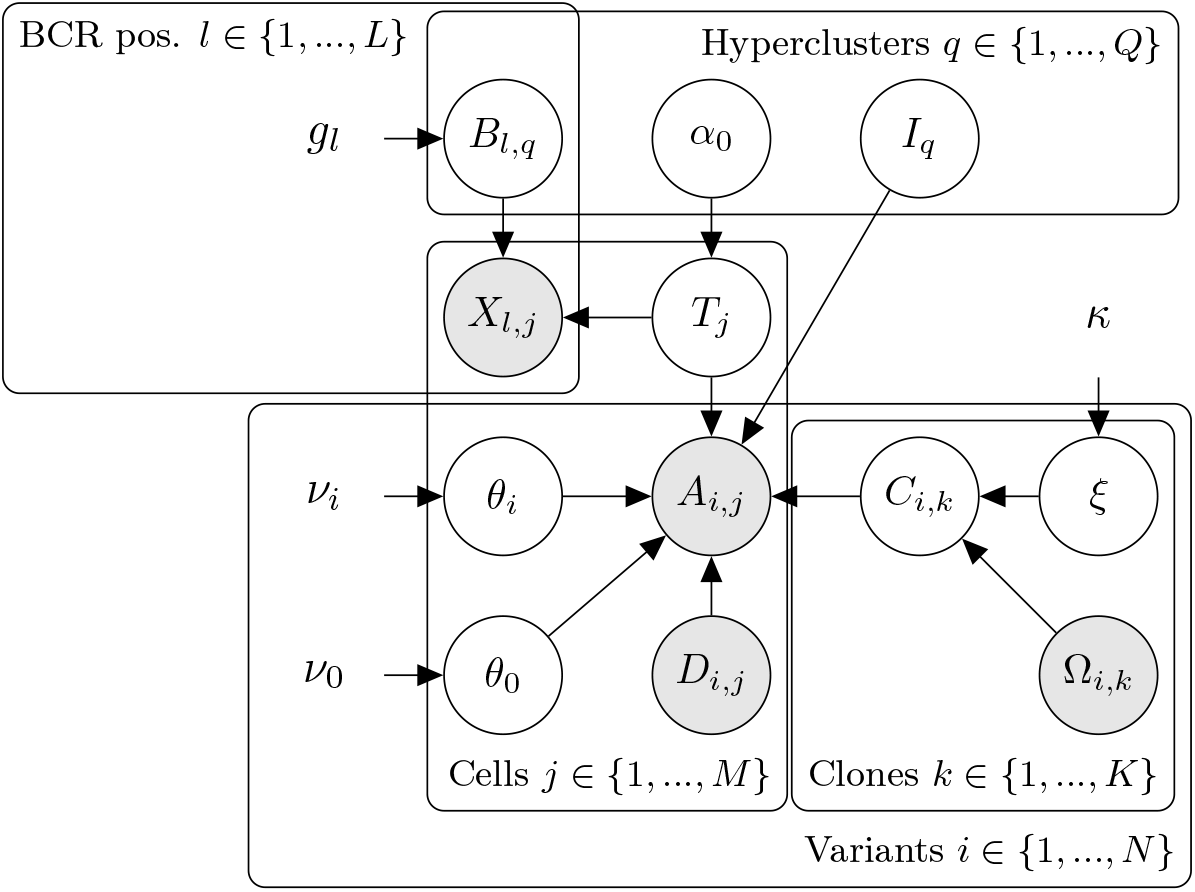
Graph of the CaClust model. White vertices represent hidden variables, grey vertices represent observed variables. Directed edges show probabilistic dependencies between variables. Vertices with no outline are model parameters. Ω_*i,k*_ is the observed genotype of clone *k* at position *i, C*_*i,k*_ being the true genotype: *C*_*i,k*_ = 1 if mutation *i* is present in clone *k* and 0 otherwise. *ξ* is the error rate between Ω_*i,k*_ and *C*_*i,k*_. *A*_*i,j*_ is the count of observed unique molecules with an alternate nucleotide at position *i* in cell *j*, with *D*_*i,j*_ being the total number of unique molecules observed at position *i* in cell *j. θ*_0_ is the probability of observing an alternate read if a cell does not carry a mutation at that position, *θ*_*i*_ is the probability of observing an alternate read if a cell does carry mutation *i. I*_*q*_ is the assignment of hypercluster *q* to one of the tumour clones. *T*_*j*_ is the assignment of a cell *j* to one of the hyperclusters and *α*_0_ is the concentration parameter of the CRP prior of *T*_*j*_. *B*_*l,q*_ is the vector of nucleotide frequencies at position *l* in BCR sequences of cells from hypercluster *q. X*_*l,j*_ is the nucleotide present at position *l* in the BCR sequence of cell *j. κ, ν*_0_, *ν*_1_, *g* are the hyperparameters of the model.

Secondly, we investigated the BCR similarity of cells in hyperclusters in K4B to hyperclusters in K5B (Fig. 5**b**). It should be noted, that no exact same BCR sequence was shared between samples; therefore, there is no immediate candidate hypercluster that could be the founder of K5B. However, we again found that cells in hyperclusters belonging to clone C3 in K4B showed the highest degree of similarity (number of BCR nucleotide mismatches) to hyperclusters in all clones from K5B, further demonstrating the evolutionary similarity of K4B clone C3 to sample K5B.

Thirdly, WES data of sample K4B contained SNVs that were not found in the WES data of sample K5B, which suggests some parallel evolution between the samples. We investigated the subclonality of 45 SNVs that were used in the K4B model inference but have not been observed in K5B WES data. For each clone, we checked how many of those 45 SNVs were inferred to be present in its genotype and how many were not. We considered a genotyping call on a variant position in a clone as reliable if the scRNA data of cells mapped to that clone contained at least one mutated read or three reference reads (Fig. 5**c**). Clone C3 has the fewest reliably called SNVs (5) not observed in K5B. This again highlights the similarity of clone C3 to the K5B sample, and while the presence of additional mutations may not make it the exact predecessor in evolution, their common ancestor could be very close up the evolution tree. This finding is in line with [14], where time-related FL samples were shown to come from a common progenitor clone (CPC), rather than being direct descendants (for divergent evolution in FL see also [31]).

We also checked the correlation of gene expression between the clones of K4B and K5B. We took the union of the top 100 most variably expressed genes in each sample and calculated the Spearmann correlation coefficient between their average expression in clones of K4B and clones of K5B (Fig. 5**d**). Here again clone C3 from K4B had the highest correlation of expression to clones from K5B; however, it has to be noted that the overall correlation scores were very high. That is expected, since it was a comparison of malignant B-cells, which had a single ancestor cell and a high overall similarity could be expected.

In summary, thanks to accurate clonal mapping using CaClust we could resolve the clonal structure and ancestry connections between the time-related samples.

## Discussion

In this work we combined in-depth molecular profiling of patient samples with probabilistic modelling to investigate the evolutionary histories and to explain the relationship between genomic and transcriptional heterogeneity in FL. To this end, we performed WES, scRNA-seq, BCR-seq and targeted resequencing and introduced CaClust, a novel method for clonal phenotype profiling with single cell genotyping in FL.

CaClust integrates BCR, WES, and scRNA information for increased accuracy and confidence. Since we consider that the evolution of BCR sequences proceeds much faster than and in parallel with the evolution of the rest of the genome, the CaClust model makes an important assumption that cells with similar BCR sequences belong to the same genetic clone. This assumption, combined with the use of nonparametric Bayesian clustering allow the new CaClust model to efficiently pool scRNA information on the clonal assignment of FL cells based on their BCR similarity. By pooling the single cells together into BCR hyperclusters, the model circumvents the biggest problem in single-cell genotyping based on scRNA sequencing data, which is the sparsity of reads in each cell. Moreover, the approach used is flexible in that it infers the optimal number of hyperclusters along with their BCR profiles from the data. As we have demonstrated both on simulated and experimental datasets, this greatly improves the clonal profile reconstruction, cell assignment, and genotyping accuracy over a rigid BCR clustering, which was previously shown to perform best in those tasks for FL [15]. The newly improved clonal and single-cell genotypes obtained with CaClust enable multiple downstream analyses that can shed light on the effects of driver mutations, possible therapeutic targets, and parallel evolution of time-related FL samples, as demonstrated on data from 4 samples from 3 patients.

However, limitations of our method still exist. Firstly, the used input scRNA data is based on 5’ end sequencing, which does not capture the full transcriptome and thus some variants could be potentially missed. Secondly, even despite the high depth sequencing and pooling effect that the hyperclusters provide for scRNA reads, some variants still did not have sufficient coverage to make reliable genotyping calls in clones. Thirdly, not all variants will result in a major phenotypic difference in their clones, thus in some cases of more homogeneous FL, the analysis will bring less discoveries.

Despite these limitations, our approach brings important insights into the ongoing debate on the sources of intratumour heterogeneity. Comparison to previous model CACTUS and simpler baselines showed that with a model that is more robust to noise and smarter in data integration, more transcriptional heterogeneity can be explained by genetic causes (Fig. 2d). Only having established the more likely genotype to transcriptional phenotype link should the remaining phenotypic variance be attributed to other effects, for which the mechanisms are less clear. With its excellent performance and rich output for downstream analysis, CaClust proved highly useful in the study of heterogeneity in FL, by extracting the phenotype-to-genotype mapping from high throughput sequencing data into an easily interpretable structure of hyperclusters and clone clusters.

## Conclusions

In this work we proposed CaClust, a novel method for clonal phenotype profiling with single cell genotyping in FL and demonstrated its potential use in the study of evolutionary histories and the relationship between genomic and transcriptional heterogeneity in FL. To our knowledge, our approach is the first to enable the joint study of these two types of heterogeneity in FL and to evaluate the strength of genotype-to-phenotype links in the evolutionary context of BCR hypermutation. Our in-depth analysis of 22492 single cells and whole exomes from four FL samples using CaClust gives insights into effects of driver mutations, possible therapeutic targets, and FL evolution.

Firstly, as model validation we showed that CaClust outperforms a state-of-the-art model on simulated and patient data. Secondly, we demonstrated that CaClust single-cell genotyping agrees with genotypes observed in an independent targeted resequencing experiment. Additionaly, our investigation of CaClust clones identified potential mutations driving clone phenotypes in patient sample K7B, which include two known pathogenic variants of *MYD88* and *CREBBP*, a *VMA21* variant causing a targetable dependency, and a novel *TSPAN33* variant; for mutations with known pathogenecity, their effects were observed in the expression phenotypes of their carriers. Lastly, the inferred clone genotypes and BCR hypercluster profiles of the time-related samples K4B and K5B gave hints of the evolutionary history of their clones, that agree with the findings on CPCs from [14].

Altogether our results illustrate that CaClust greatly facilitates an effective study of the extensive genomic and transcriptional heterogeneity in FL and their link, by providing the first method for their joint analysis utilising the context of BCR hypermutation.

## Methods

### Data collection

#### Patient sample collection

Samples with histologically confirmed infiltration of follicular lymphoma (FL) grade 1-2 were collected according to the Declaration of Helsinki and under authorization of applicable biobank regulations of Leiden University Medical Center and the Ethical Committee of Leiden University Medical Center (reference HEM 008/SH/sh). Excisional lymph node biopsies were processed immediately by gentle mechanical disruption and mesh filtration. Single cell suspensions were frozen in 10% DMSO and remaining tissue fractions were cultured in low-glucose (1g/L) DMEM with 8% fetal bovine serum to obtain adherent cell cultures for isolation of DNA of cells representing normal counterpart.

#### Cell processing, library preparation and sequencing

FL cells were thawed and purified by flowcytometry using anti-CD19-APC (Becton Dickinson, Franklin Lakes, NJ) and anti-CD10-PECy7 (Becton Dickinson) followed by removal of dead cells (MACS Dead Cell Removal Kit, Miltenyi Biotech, Bergisch Gladbach, Germany). For whole exome sequencing, DNA was isolated from 1 · 10^6^ purified FL cells and from 0.5 · 10^6^ cultured adherent cells (Allprep DNA/RNA Mini Kit, Qiagen, Hilden, Germany). Whole exomes were sequenced using SureSelect Human All Exon V7 baits (Agilent, Santa Clara, CA). Adherent cells representing normal cells were sequenced at 50x coverage. To discriminate between early clonal variants and more recently acquired subclonal variants as putative drivers of distinct clones, FL bulk DNA was sequenced at 1500x coverage to allow reliable calling of rare variants down to a variant allele frequency (VAF) of 0.02. For 5’ based single cell transcriptome sequencing, 1 · 10^5^ similarly purified viable cells were loaded on a Chromium X single cell device to generate cDNA libraries for an expected 6 · 10^3^ − 8 · 10^3^ cells per sample. (10X Genomics, San Fransisco, CA) Inside the 10X Genomics chip, single cells and oligonucleotide-covered beads are simultaneously captured as aqueous droplets in oil. Per bead, all oligonucleotides share an identical single cell barcode (scbc), and every single oligonucleotide molecule carries a unique molecular identifier (UMI). In the droplet, cells are lysed and cDNA synthesis is 3’ primed with oligo-dT. After amplification of the primary cDNA library by using universal primers, the amplified library is split in 3 fractions for 1) full transcriptome sequencing, 2) enrichment of BCR transcripts followed by full length sequencing, 3) targeted resequencing of subclonal somatic variants. Sequencing for WES and single cells was performed on HiSeq2500 or HiSeq4000 devices (Illumina, San Diego, CA).

#### Variant calling

FASTQ files from whole exome sequencing (WES) were processed using the Sarek workflow v2.7 and aligned to the human reference genome GRCh38 using Burrows Wheeler Algorithm (BWA) v0.7.17. [32, 33] Duplicated mapped reads were marked, local realignment of regions flanking indels and recalibration of base quality scores were performed to obtain more accurate bases according to the Genome Analysis ToolKit (GATK) best practices version v4.1.7.0. [34] Single nucleotide variants (SNV) and short insertions and deletions (INDELS) were called using Strelka2 v2.9.10. [35] Only high confidence variants defined by quality scores (GQX) of at least 15 for SNV and 30 for INDELS were kept. Pathogenecity of variants was determined using the Geneticist Assistant NGS Interpretive Workbench (SoftGenetics) based on publicly available variant-databases (dbSNP, ClinVar, and COSMIC) and literature, into class 1 (benign), class 2 (likely benign), class 3 (unknown significance), class 4 (likely pathogenic), or class 5 (pathogenic) [36–38].

#### Variant selection for CaClust modeling

We chose only somatic single nucleotide variants called from Strelka that could also be observed in the scRNA data of the cells with complete BCR heavy and light chain sequences, and that showed at least one alternate read across the cells.

Some germline variants were present in the Strelka output, as they had an increased frequency of the alternate nucleotide over the normal sample. We chose to include only those variants that showed over 3-fold increase in the frequency of the alternate nucleotide between the normal and tumour sample. The full table of included variants per sample can be seen in Supplementary Data 1.

### Copy number inference (FalconX)

We use FalconX [39] for the inference of copy number alteration (CNA) events. In the getASCN.x method we use a threshold of 0.1; later, in the quality-filtering falconx.qc we set the CNA length.threshold of 10^7^ basepairs and the delta copynumber threshold of 0.1.

### Inference of input clonal profiles

We use CANOPY [40] for the estimation of input clonal profiles for the model. We run 5 CANOPY chains for each clone number *K* ∈ 3, *…*, 6, choosing the number of clones for each sample with the highest BIC score. The minimum number of model iterations is set to 20000 and the maximum to 100000. The input CNAs for CANOPY inference are obtained with FalconX.

### The CaClust model formulation

CaClust can be seen as a significant extension to our previous model, CACTUS, with the functionality of non-parametric Bayesian clustering applied to BCR sequences.

We assume we are given a cancer tissue sample with WES, scRNA and BCR receptor profiling. Let *i* ∈ {1, …, *N*} denote a position of a SNV that can be found both in WES and scRNA data. We assume that a set of *k* ∈ {1, …, *K*} clones is given, each with a distinct genotype. We describe the given clone genotypes with a matrix **Ω**, where an entry Ω_*i,k*_ is 1 if variant *i* is present in the given genotype of clone *k* and 0 if it is not.

As in CACTUS, the input matrix of clone genotypes is assumed to be imperfect, containing errors with rate *ξ*. We take *ξ* with a prior distribution Beta with parameters *κ* = (*κ*_0_, *κ*_1_), obtaining ℙ (*ξ*|*κ*) = Beta(*ξ*; *κ*_0_, *κ*_1_). We then introduce the matrix **C**, where *C*_*i,k*_ are the hidden variables representing the true genotype of clone *k* at variant position *i*, such that:

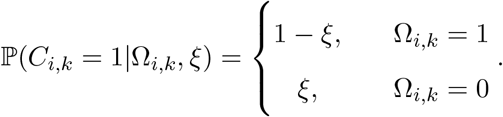

The main advantage of CaClust over its predecessor CACTUS is the way the clustering of cells by their BCR receptors is modelled. Let *j* ∈ {1, …, *M*} be the cell indices. *T*_*j*_ denotes the BCR hypercluster to which cell *j* is assigned. Contrary to its predecessor, CaClust does not consider a fixed number of BCR hyperclusters, but rather allows it to be inferred, using the Chinese Restaurant Process (CRP) for the prior, i.e.:

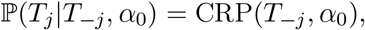

where *T*_−*j*_ is the hypercluster assignment of all cells but *j* to the BCR hyperclusters in the model, and *α*_0_ is the concentration parameter.

We characterise each BCR hypercluster *q* of cells with its BCR frequency profile *B*_*q*_ and with **B** we denote all those profiles in the model. Let *l* ∈ {1, …, *L*} index BCR positions. Then the vector *B*_*l,q*_ = [*B*_*A,l,q*_, *B*_*C,l,q*_, *B*_*G,l,q*_, *B*_*T,l,q*_] describes the probabilities, with which a cell belonging to hypercluster *q* has at position *l* a nucleotide *A, C, G* or *T*, respectively. We set a Dirichlet prior on *B*_*l,q*_ with parameters *g* = (*g*_*A*_, *g*_*C*_, *g*_*G*_, *g*_*T*_):

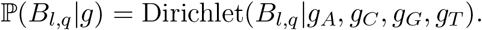

For each cell *j* we are given its BCR sequence as *X*_*j*_, where *X*_*j,l*_ is the nucleotide at position *l*; with **X** we denote the matrix of all cells’ BCR sequences. Given the hyperclustering and its BCR frequency profiles we treat each cell’s BCR as coming from a categorical distribution with probabilities described by its hypercluster’s profile. So for a cell *j*, its hypercluster *T*_*j*_ and the BCR frequency profile we have:

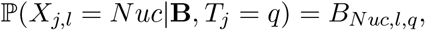

where *Nuc* is one of the four nucleotides, *Nuc* ∈ {*A, C, G, T*}.

We make the assumption that cells from the same hypercluster belong to the same tumour clone, so we want to find the hypercluster to clone assignment. By *I*_*q*_ we denote the tumour clone that hypercluster *q* is assigned to and with **I** the assignment of all hyperclusters in the model. We make no prior assumptions on that assignment and so we set a uniform prior distribution: 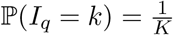.

From the scRNA data we create matrices **A** and **D**, where *A*_*i,j*_ and *D*_*i,j*_ are the numbers of alternate and total reads respectively, that map to position *i* in cell *j*. We then define observation probabilities *θ* = (*θ*_0_, *θ*_*i*_): *θ*_0_ is the probability of a read being mutated if it comes from a cell mapping to a clone that does not have that mutation in its genotype, *θ*_*i*_ is the probability of a read being mutated if it comes from a cell mapping to a clone that does have mutation *i* in its genotype. Then, the likelihood of observing *A*_*i,j*_ mutated reads from *D*_*i,j*_ total reads is:

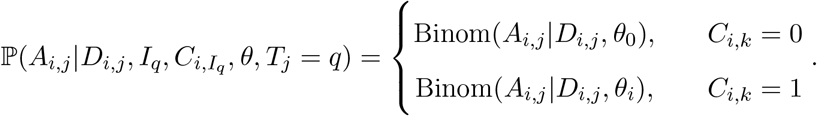

We pick beta priors for *θ*_0_ and *θ*_*i*_, with parameters (*a*_0_, *b*_0_) and (*a*_1_, *b*_1_) respectively:

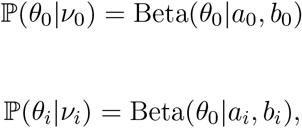

where we denote *ν*_0_ = (*a*_0_, *b*_0_), *ν*_*i*_ = (*a*_*i*_, *b*_*i*_) and *ν* = (*ν*_0_, *ν*_1_).

Let *A*_*q*_ = {*A*_*i,j*_}_*j*∈*q*_, *D*_*q*_ = {*D*_*i,j*_}_*j*∈*q*_ be scRNA reads from cells in hypercluster *q*. Since we assume that scRNA reads at different positions or from different cells are conditionally independent, then the total likelihood of these reads is:

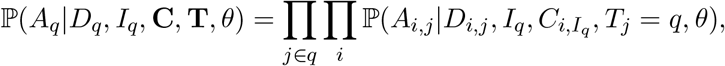

### Gibbs sampling

Inference in CaClust is performed using a Gibbs sampler, where each variable is iteratively sampled from its conditional probability given the current values of the other variables in the model. Since CaClust is a probabilistic graphical model, this conditional probability is equivalent to the conditional probability of the variable given its Markov Blanket [41]. The sampling of variables related to the CRP is performed using a dedicated procedure described below. The inference is carried out until convergence, as measured by the Gelman-Rubin diagnostic; afterwards, the samples from iterations after burn-in approximate the true posterior distribution of the variables.

### Conditional probabilities of variables

In the Gibbs sampler, we sample the variables from their conditional probabilities given their Markov Blankets (MB). Using Bayes’ rule we factor these probabilities as follows.

For the error rate *ξ* we have:

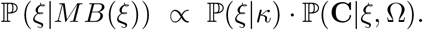

The prior on *ξ* is a Beta distribution with parameters (*κ*_0_, *κ*_1_) and the likelihood ℙ(**C**|*ξ*, Ω) is a product of Binomial distribution functions over variables *C*_*i,k*_ for *i* ∈ 1, *… N* and *k* ∈ 1, *… K*, where a success is defined as a disagreement (since *ξ* is an error rate) between Ω _*i,k*_ and *C*_*i,k*_. Therefore, from the Beta-Binomial conjugacy we obtain:

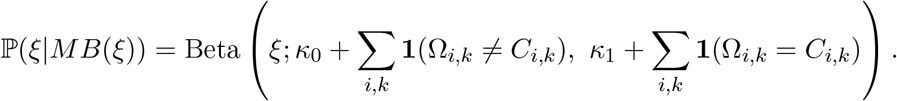

For the true genotypes *C*_*i,k*_ we have:

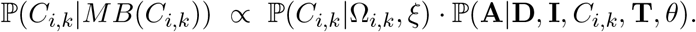

Since we assume reads at different variant positions *i* and reads in different cells *j* are conditionally independent, the above probability factorises as:

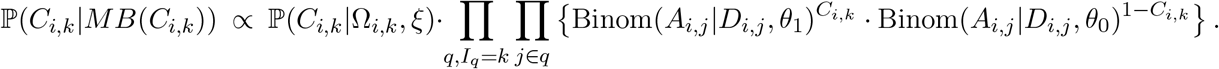

For the hypercluster-clone assignment variable *I*_*q*_ we have:

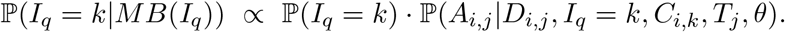

We use an uninformative prior on *I*_*q*_, so the posterior probabilities of *I*_*q*_ are proportional to the likelihoods of scRNA reads, which again we assume to be conditionally independent across positions *i* and cells *j*:

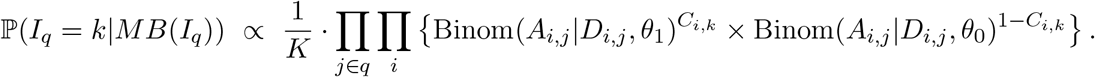

For the hyperclustering variable *T*_*j*_ we have:

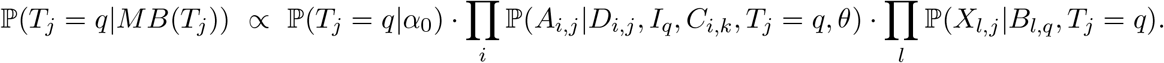

The observations of BCR sequences across positions *l* and cells *j* are also conditionally independent, so the above factorises to:

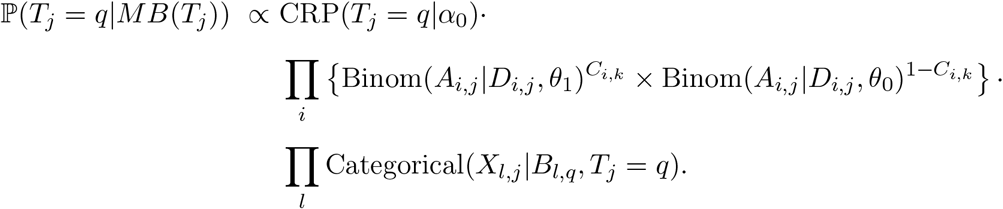

We can calculate the above for all hyperclusters *q* that are non-empty. However, since we are dealing with a CRP, then during the sampling of cell *j*’s hypercluster assignment *T*_*j*_ we need to include the possibility of joining a new hypercluster. So during the sampling of *T*_*j*_, if *Q* is the number of non-empty hyperclusters in clustering *T*_−*j*_, we add a hypercluster *Q* + 1 with parameters *I*_*Q*+1_, *B*_*Q*+1_ sampled from their prior distributions. Then we can calculate the above for hyperclusters 1, *…, Q* + 1 and sample *T*_*j*_ with appropriate probabilities.

For the hypercluster BCR frequency profiles *B*_*l,q*_ we have:

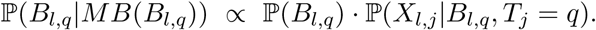

Since the *B*_*l,q*_ variable has a Dirichlet prior and the observations of nucleotides *X*_*l,j*_ come from a categorical distribution with probabilities *B*_*l,q*_, then from the Dirichlet-Multinomial conjugacy we get:

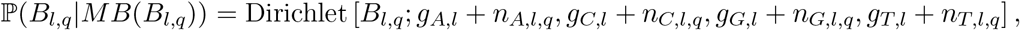

where *n*_*Nuc,l,q*_ is the number of occurances of nucleotide *Nuc* at position *l* in BCR sequences of cells belonging to hypercluster *q*.

For the variant read observation probabilities *θ* we have:

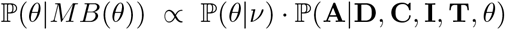

Since *θ*_0_ and *θ*_*i*_ have a Beta prior and the likelihoods of **A** are Binomial distributions with sizes **D**, then from the Beta-Binomial conjugacy we get:

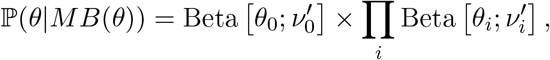

where 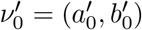 and 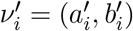 are defined by:

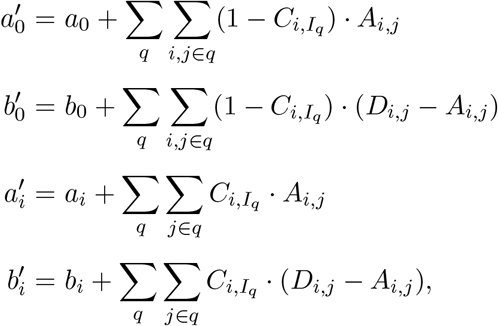

The conditional probability of *α*_0_ is a special case and its sampling is described in the following section.

### Updating the concentration parameter *α*_0_

During model inference we also sample *α*_0_, the concentration parameter of the Chinese Restaurant Process behind our clustering. For sampling *α*_0_ we use a method described in paper [42]. It applies to mixture models in general, and in this section we explain how it is implemented in the CaClust model.

Firstly, we assume a Gamma prior over *α*_0_,

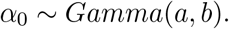

We have *M* as the number of cells, and let *Q* be the number of non-empty hyperclusters in current sampling iteration. Since *Q* is not fixed, we consider *Q* as another random varable. In the Chinese Restaurant Process (CRP) we have:

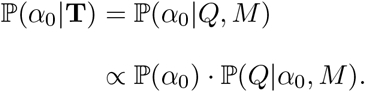

Then from the probability density of CRP we have:

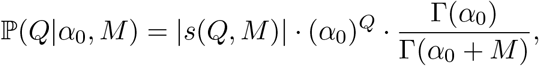

where *s*(*Q, M*) are Stirling numbers of the first kind and, more importantly, they are independent from *α*_0_. For *α*_0_ *>* 0 (as in our case), we can rewrite the gamma functions from above as:

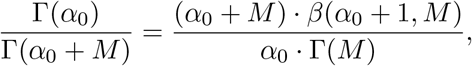

where *β* is the beta function. So, since Γ(*M*) is constant w.r.t. *α*_0_, the conditional probability of *α*_0_ takes the form:

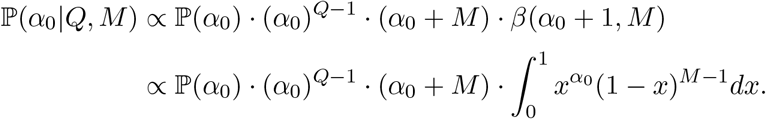

This shows that ℙ(*α*_0_|*Q, M*) is a marginal probability distribution of a joint distribution of pairs (*α*_0_, *σ*), where:

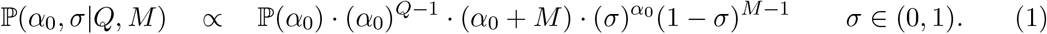

With the *Gamma*(*a, b*) prior on *α*_0_ we obtain the following probabilities:

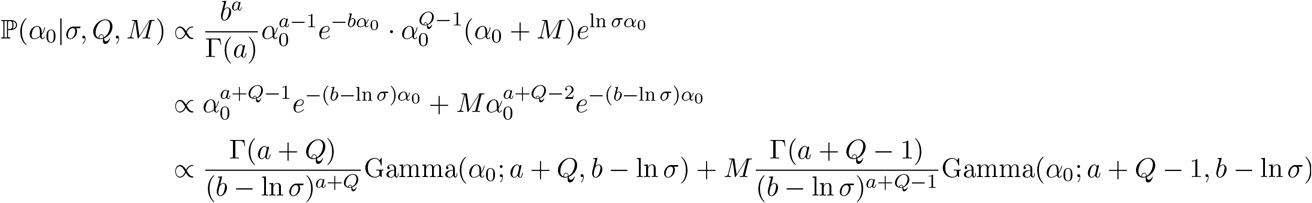

which is a mixture of two gamma densities:

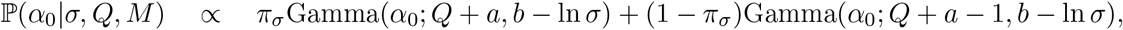

with weights such that 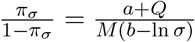.

Lastly, by marginalising Eq.1 w.r.t. *α*_0_, we get the conditional distribution of *σ*:

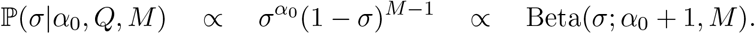

Therefore, we sample *α*_0_ at each iteration as follows:

1. pick a new value for *σ* with previous values of *Q* and *α*_0_
2. pick a new value for *α*_0_ with previous value of *Q* and new *σ*.

### Data simulation methods

In this section we describe the process of simulating data for evaluation of model performance, specifying the parameter settings for different simulation runs and evaluation metics.

### Simulation process

The input data required by the model for inference are: matrix **D** of read counts over mutations in the cells, matrix **A** of alternate read counts over mutations in the cells, BCR sequences of cells, and matrix **Ω** of observed clone genotypes.

Assume we are given: the number of clones *K*, the number of cells *M*, the number of variant positions *N*, the length of BCR sequences *L*. Then the simulation steps are as follows:

- Step 1: Hyperclustering simulation.
  1. Generate the concentration parameter *α*_0_ from its prior distribution.
  2. Using the CRP with concentration parameter *α*_0_ cluster the *M* cells.
  3. Assign each of the *Q* resulting hyperclusters to one of the *K* clones with uniform probability.

- Step 2: BCR sequence (**X**) simulation.
  1. For each hypercluster *q* and each BCR position *l*, sample its BCR frequency profile *B*_*l,q*_ from the Dirichlet distribution with prior parameters *g*_*l*_.
  2. For each cell *j* simulate its BCR sequence at position *l* from a categorical distribution with the frequencies *B*_*l,q*_ that we obtain for its hypercluster *q* from the previous step. In simulation scenarios with centroid behaviour (see Simulation types) we additionally randomly select hyperclusters with rate *r*_*clust*_ and within them *r*_*cell*_ cells to express identical BCR sequences, equal to the sequence that is most probable given their hypercluster’s profile *B*_*l,q*_.

- Step 3: **Ω** and **C** simulation.
  1. Simulate *θ*_0_, *θ*_*i*_, *ξ* from their prior distributions with desired parameters.
  2. Pick a variant rate *v*.
  3. For each clone simulate its mutation profile *C*_*k*_: at each variant position *i* we have a chance of *v* that clone *k* exhibits a variant at that position i.e. *C*_*i,k*_ = 1.
  4. Simulate the observed matrix **Ω** by randomising **C** with error rate *ξ*.

- Step 4: scRNA-seq variant data (**A** and **D**) simulation:
  1. For each cell *j* and variant position *i* sample the total number of observed reads mapping to that position (*D*_*i,j*_) from a Poisson distribution with mean *μ*_*D*_.
  2. For each cell *j* and each position *i* sample the number of observed variant reads *A*_*i,j*_ from its conditional probability distribution.

### Choosing simulation parameters

In the simulation process we have control over the values of several parameters, which influence the simulated structure in different ways. We can divide those parameters into three groups: data dimensions, simulation type agnostic, and simulation type specific.

### Data dimension parameters

Data dimension parameters are fixed for all simulation types and affect primarily the computational times.

We set the number of tumour clones *K* = 3, the number of variant positions *N* = 100 in the tumour genotypes (fixed according to the numbers of variant positions after filtering in the analyzed real patient data), the number of cells *M* = 1000. Finally, we set the number of mutated BCR positions *L* = 300, since in the experimental data we observed between 200 and 400 positions that contain at least one alternate nucleotide. Note that this number is smaller than the combined length of the BCR heavy and light chain sequences, which is around 600-700 nucleotides.

### Simulation scenario-agnostic parameters

These parameters are shared between all simulation scenarios. Firstly, the variant rate *v* used to generate the clone genotypes; for each clone *k* and each variant position *i* we set a probability *v* = 0.3 that *C*_*i,k*_ = 1. Secondly, the parameters *ν* = (*ν*_0_, *ν*_1_) of error rate *ξ*’s beta prior distribution; we set *ν*_0_ = 1, *ν*_1_ = 19.

Lastly, the prior parameters of alternate nucleotide observation probabilities *θ*_0_ and *θ*_*i*_. In the model we use only somatic variants, so *θ*_0_ reflects the probability that an alternate nucleotide from a different position was mismapped. We assume high quality of position analysis, thus *θ*_0_should be low. Secondly, we assume we are dealing with heterozygotic somatic variants; therefore, the probability of observing a variant nucleotide from the mutated allele is on average 50%, but can vary due to the random nature of mRNA expression and sequencing. To reflect the above, we choose the prior parameters of *θ*_0_ to be *a*_0_ = 0.2, *b*_0_ = 99.8, and the prior parameters of *θ*_*i*_ to be *a*_*i*_ = 4.5, *b*_*i*_ = 5.5.

### Simulation scenarios

We wanted to test model performance on datasets with three varying characteristics: scRNA read depth, controlled with *μ*_*D*_; number of BCR hyperclusters, controlled with *α*_0_; and intracluster variance of BCR sequences, controlled with *s*_*g*_, *r*_*clust*_, *r*_*cell*_. To do this, we created a base scenario modelling medium read depth (*μ*_*D*_ = 0.01), low number of BCR hyperclusters (*α*_0_ = 5), and medium intracluster BCR variance (*s*_*g*_ = 0.01, *r*_*clust*_ = 0, *r*_*cell*_ = 0). Then, in each simulation type we changed one of these parameters, while keeping the rest at base values.

Apart from the base scenario we created eight following simulation scenarios. Two scRNA read depth scenarios with high, or low average numbers of reads per variant in a cell; these scenarios reflect a high and low quality sequencing experiments with high and low scRNA information respectively. One sparse hyperclustering scenario with numerous BCR hyperclusters in the data; this models a tissue with multiple BCR clusters evolving in parallel. A scenario with high variance BCR sequences within hyperclusters; this models cells with highly mutated BCR sequences that give low information. And lastly, four scenarios with *x*% of cells within *y*% of hyperclusters sharing their hyperclusters’ prevalent BCR sequence (with *x, y* ∈ {20, 80}); this accounts for BCR hypercluster tendencies observed in real data, in which most hyperclusters contain a large subset of cells with identical BCR sequences. All simulation types are shown in Table 1 along with the values of the varied parameters.

**Table 1:** Simulation scenarios with their specific parameters. Parameters in blank spaces are the same as in the basic scenario.

| Simulation scenario | $\mu_D$ | $\alpha_0$ | $s_g$ | $r_{clust}$ | $r_{cell}$ |
| --- | --- | --- | --- | --- | --- |
| Basic | 0.01 | 5 | 0.01 | 0 | 0 |
| High reads | 0.1 |  |  |  |  |
| Low reads | 0.001 |  |  |  |  |
| Sparse hyperclustering |  | 50 |  |  |  |
| High variance BCR |  |  | 0.1 |  |  |
| 80% centroids in 80% of hyperclusters |  |  |  | 0.8 | 0.8 |
| 80% centroids in 20% of hyperclusters |  |  |  | 0.8 | 0.2 |
| 20% centroids in 80% of hyperclusters |  |  |  | 0.2 | 0.8 |
| 20% centroids in 20% of hyperclusters |  |  |  | 0.2 | 0.2 |

**Table 2:**
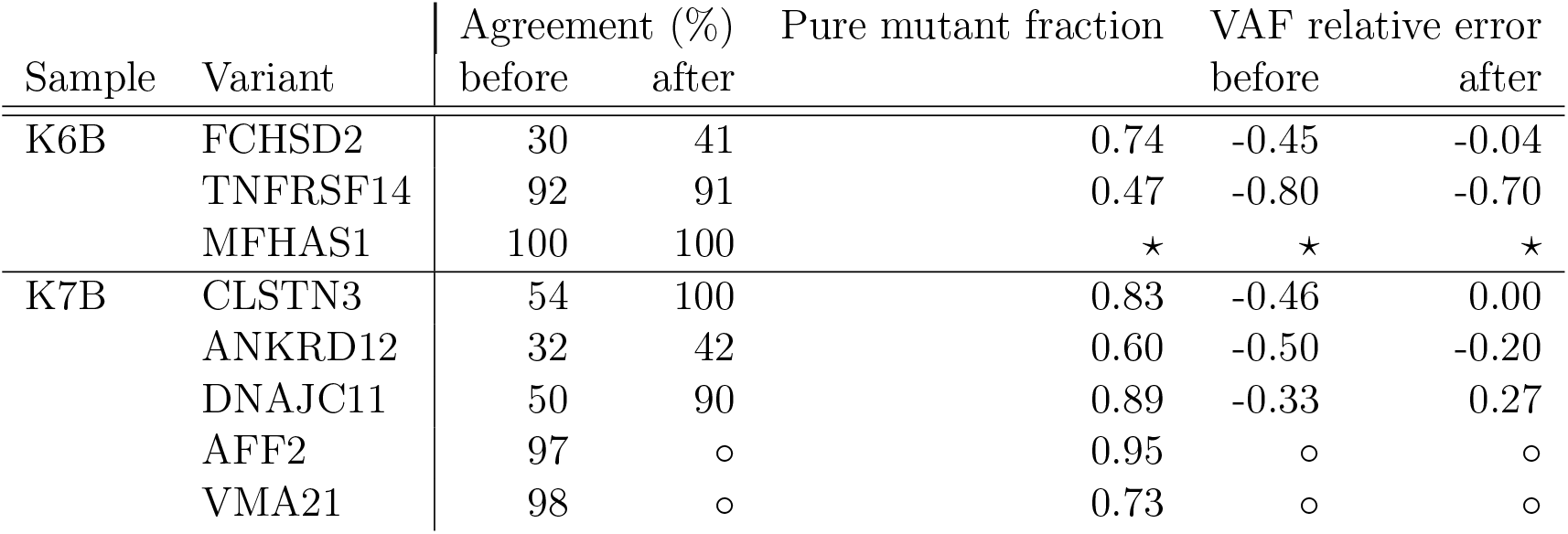
Agreement of CaClust and resequencing results before and after the correction for random monoallelic expression. Pure mutant fraction is the ratio of cells with only mutant UMIs to cells with any mutant UMIS in the resequencing. VAF relative error is calculated as the relative difference between the true VAF from WES and the VAF predicted with resequencing genotypes before or after the correction. ⋆ – no variant *MFHAS1* reads were detected in resequencing; ∘ – the correction does not apply to monoallelic *AFF2* and *VMA21*

### Performance metrics

To measure the performance of CaClust and CACTUS on simulated datasets we define three performance metrics: cell to clone assignment accuracy, genotype reconstruction accuracy, and hyperclustering reconstruction agreement.

We calculate the cell to clone assignment accuracy as the fraction of cells that were assigned to their correct clone in the MLE assignment after model inference.

Secondly, the genotype reconstruction accuracy measures how well the model corrects the input matrix of clonal profiles **Ω**, which contains errors with rate *ξ*. To calculate it, we take the MLE of the true clone genotypes **C** from the model and compute the fraction of entries that it agrees on with the hidden clonal profiles from the simulation.

Finally, with the hyperclustering reconstruction agreement we measure the similarity between the hyperclustering from the data generation process and the hyperclustering reconstructed by the model. For this, we calculate the adjusted Rand index between the final hyperclustering of cells in the model and the hyperclusters from the data generation process.

### Model application process

#### Inference parameters

For model inference we need to choose the following prior parameters.

Firstly, the parameters of the beta priors of *θ*_0_, *θ*_*i*_, which are *ν*_0_ = (*a*_0_, *b*_0_), *ν*_*i*_ = (*a*_*i*_, *b*_*i*_). Since *θ*_0_ should be the small probability of an observed read being variant in a clone not containing that mutation in the genotype, we set the prior parameters *a*_0_, *b*_0_ such that 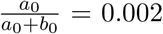 and that their magnitude *a*_0_ + *b*_0_ is equal to the number of total reference reads in the sample. *θ*_*i*_ is the probability of an observed read at position of SNV *i* being variant in a clone containing SNV *i*. To account for bursty expression and harder mapping of variant fragments we set *a*_*i*_, *b*_*i*_ such that 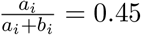 and their magnitude *a*_*i*_ + *b*_*i*_ to be equal the total number of variant reads of SNV *i* in the cells.

Secondly, the prior parameters of the BCR frequency profiles. At each BCR position we use a low strength uninformative prior of *g* = (0.01, 0.01, 0.01, 0.01), which influences more data-driven frequency profiles.

For the beta prior on the input genotype error rate *ξ*, we use parameters *κ*_0_, *κ*_1_ with values such that 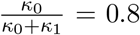 and their magnitude *κ*_0_ + *κ*_1_ = *N · K*, where *N* is the number of SNVs and *K* is the number of clones used for the sample. In that way the genotype at each clonal position is a priori weighed *κ*_0_ : *κ*_1_ in favour of the input genotype call; during the inference that ratio changes with the numbers of agreements and disagreements between **Ω** and **C** in the model, which are on the level of *N · K*.

For the gamma prior of *α*_0_ we use a non-informative prior with parameters (1, 1).

#### Model initialisation

We initialise the hyperclustering in the model (*T* variable) with the clustering of cells with identical BCR sequences. This is to promote faster convergence and is in line with the assumption, that hyperclusters of cells with identical BCR sequences should come from the same evolutionary clone. The model can also be initialised with random hyperclusters or hyperclusters containing singular cells.

After hypercluster initialisation we obtain the initial assignment of hyperclusters to clones by performing initial Gibbs sampling iterations only for **I, C**, *ξ, θ*_0_, *θ*_*i*_ variables, while keeping the initial BCR hyperclustering constant. This is done to ensure that before relaxing the BCR hyperclustering in full model interations we resolve any major disagreement between the input clonal profiles and the scRNA variants observed in the initial BCR hyperclustering.

Afterwards, the *α*_0_ and **B** variables are initialised from their conditional probabilities and the full model sampling iterations are ready to be performed.

#### Convergence assessment

The model is run for a set number of initial and full sampling iterations; then, for assessing the model convergence we use the Gelman-Rubin diagnostic with stable variance estimators [43] on the variables with continous values in the model (*θ*_0_, *θ*_*i*_, *α*_0_). The model is assumed to have converged if between the chains in a sample we observe the multivariate potential scale reduction factor (MPSRF) lower than 1.001. If not converged, the next set of full sampling iterations is carried out and the convergence is reassessed. We use 5000 initial and 500 full sampling iterations.

#### Result extraction

For each sample we choose the model chain with the highest likelihood; then, from that chain we take the maximum a posteriori (MAP) of cell-clone assignment and clone genotypes, and the output hyperclusters are taken to be the hyperclustering with maximum likelihood through the iterations.

#### Independent gene expression clustering

Gene expression clusters were obtained by PCA dimensionality reduction to 30 dimensions and next Leiden clustering on the reduced profiles with resolution parameter of 0.3m using Seurat package [44].

#### Pathogenecity prediction

All variants were annotated within the Geneticist Assistant NGS Interpretive Workbench (Soft-Genetics) by public variant-databases (dbSNP, ClinVar, and COSMIC) and available literature, into class 1 (not pathogenic), class 2 (potentially not pathogenic), class 3 (unknown significance), class 4 (potentially pathogenic), or class 5 (pathogenic). [36–38]

#### Differential gene expression analysis

We use the DESeq2 package [45] for differential gene expression analysis in the samples. For each clone we use its cells as a test group and the rest of the cells as a reference group. The analysis is carried out on the SCT counts, with size factors estimated using the scran package [46]. For significance we use the likelihood-ratio test (LRT) implemented in the package, with Benjamini-Hochberg correction for multiple testing.

From the DE analysis we exclude: genes with a total read count *<* 1% of the cell count in the data; the ribosomal protein L and S genes; the immunoglobulin IG[HKL] genes; and the mitochondrial MT-genes.

#### Gene set enrichment analysis

We perform Gene set enrichment analysis (GSEA) as described in [47] on the 50 hallmark genesets [48]. For each clone we use as input the list of genes ranked by their foldchange found in the DE analysis. For FDR in the GSEA we use gene label reshuffling with 10000 permutations.

### Targeted resequencing

#### Variant selection for targeted resequencing

Variants with subclonal VAF (0.05<VAF<0.4 or 0.6<VAF<0.8) in genes that were detectable in ≥ 2.5% of single cells based on single cell transcriptome data were selected. Variants that were synonymous, located outside exons, in immunoglobulin genes or more than 2000 bp from start of transcripts were excluded.

#### Targeted resequencing procedure

Pools of 10X Genomics cDNA that were remaining after GEX and BCR sequencing were used as template for targeted resequencing. A semi-nested 2-step amplification strategy was designed based on a reverse outer and a reverse nested inner primer both located 3’ of the target variant position (see Supplementary Data 2). For primer validation, an additional forward primer in the 5’ region of the gene was designed and used on cDNA that was generated from bulk-sorted FL cells as described previously with modifications: initiation of reverse transcription using oligo-dT and 5’extension with an alternative 5’ template switching oligo [49]. Validation PCRs were performed with 10 μL bulk FL derived cDNA template using Phusion Flash High Fidelity PCR Master Mix (Thermo Fisher Scientific, Waltham, MA) with the reverse outer gene specific primers and enrichment primer I both at 1 μM. As PCR program was used: melting 45 sec 98°C, amplification for 20 cycles: melting 20 sec 98°C, annealing 30 sec 67°C, elongation 120 sec 72°C, followed by a final elongation step of 120 sec 67°C. Aliquots of 10 μL PCR product were run on 1% agarose gel and visualized. If bands were visible, remaining 40 μL PCR products were purified using AMPure XP Beads (BeckmanCoulter, Indianapolis, IN). The nested PCR was performed under identical conditions with the inner gene specific primer and enrichment primer II. If no bands were visible, alternative primers were designed and tested. Using validated primer sets, aliquots of 1.5 μL of 10X Genomics single cell cDNA pools were amplified with enrichment primers I and II and the validated outer and inner 3’ reverse primers under identical conditions as in primer validation. After the second amplification, PCR products were run on preparative 1% agarose gel, visible bands were excised, purified using Promega Wizard PCR Preps DNA Purification System (Thermo Fisher Scientific) and run on Bioanalyzer (Agilent) for accurate quantification of obtained PCR products. Equimolar amplicon pools were generated and single molecule sequencing was performed on PacBio Sequel II platform (PacBio, San Diego, CA).

#### PacBio full length sequence data processing

Circular consensus sequence fastq files were used as input for filtering and genotype calling. PacBio polymerizes circularized single strand DNA templates and dependent on the start of the reaction results in either the forward or reverse sequence. To obtain all reads in the forward direction, a copy of every read was reverse complemented and added to the data. Using the 5’ end PCR adapter sequence AATGATacg, also allowing deletion of 1-3 nt thus accepting aATGATAcg, aaTGATACg and aatGATACG, were used to keep only reads in the forward direction. For reads that were not correctly split during circular consensus generation and thus consisted of the forward linked to the reverse sequence, mean quality score of nt 5-100 of each sequence end was calculated. The read part with lower quality was clipped. Next, alignment scores were obtained for read nucleotides (nt) 47 to 60 with the distal PCR adapter motif CTTCCGATCT, and read nt 82 to 100 with TSO+G motif TTTCTTATATG. The space between adapter and TSO comprises the single cell barcode (scbc) of 16 nt and the unique molecular identifier (umi) of 10 nt, and thus should be 26 nt. Reads with adapter motif alignment score ≥ 9 and, TSO+G alignment score ≥ 10 and a scbc-umi space of 24 to 28 nt were accepted. The latter filter taking into account potential insertions and deletions within polyhomologous stretches. In the second step, reads were aligned with wildtype and mutant reference sequences of 41 nt with 20 up- and downstream nt flanking the nt substitution or insertion/deletion. Reads that aligned with a single reference with alignment score ≥ 30 of max 41 and nt quality score ≥ 120 of max 126 at the variant position were accepted. In the third step, reads were assigned to cells using as reference scbc from valid cells from CellRanger default output for gene expression profiling and BCR VDJ/VJ sequencing. Nt from end of adapter −1 to +18, thus a stetch of 19 nt was aligned with all reference scbc. Matches with a single scbc and alignment score ≥ 14 and an average quality score of ≥ 110 of max 126 were accepted. Two potential umis were extracted, one of exactly 10 nt downstream of the scbc (umi10), and the second one between the end of the scbc until the last nt before the TSO start (umiTso). No reference for umis is available, and we expect due to amplification and sequencing errors an overestimation of the repertoire of umis. We therefore collapsed highly similar umis as follows: Per scbc and gene, identical umis were counted and arranged by decreasing count. With the most dominant umi as reference, for all other umis a pairwise alignment score was calculated. A discrepancy of max 3 nt was allowed for a umi to be collapsed into the more dominant umi. Independently for umi10 and umiTso, the process was iterated over all umis. If the resulting umi of both umi10 and umiTso was identical, collapsed umis were corrected towards the dominant umi. If the resulting umi of umi10 and umiTso were not identical but their Levenshtein distance was ≤ 3, umi10 was chosen as the final umi. At Levenshtein distance >3, the read was rejected. Genotyping of umis and scbc was performed as follows: In case more than 1 umi was detected per single cell and gene, we counted the number of duplicates and evaluated if all umis had the same genotype. If all reads had identical genotypes, the umi genotype was called accordingly. As a result of amplification errors however, chimeric PCR products can be formed resulting in mixed wildtype and mutant reads withing 1 umi. Umi genotypes were called based if mutant or wildtype read counts were at least 0.67 of the total umi read count. Single cells were called mutant if at least 1 mutant umi was detected. Wildtype was called only if no mutant umis and at least 4 wildtype umis were detected in the absence of mutant umis.

## Declarations

### Ethics approval and consent to participate

Samples with histologically confirmed infiltration of follicular lymphoma (FL) grade 1-2 were collected according to the Declaration of Helsinki and under authorization of applicable biobank regulations of Leiden University Medical Center and the Ethical Committee of Leiden University Medical Center (reference HEM 008/SH/sh).

### Consent for publication

Not applicable.

### Availability of data and materials

Single cell gene expression and B-cell receptor transcriptome data is available in NCBI GEO repository identifiers GSE252687, GSE252642, GSE252416 and GSE252344B. Whole exome sequencing data is available in NCBI Sequence Research Archive BioProject id PRJNA1062119.

CaClust model code and input data available at github.com/szczurek-lab/CaClust.

## Authors’ contributions

K.O.O., S.D.S., C.A.M.v.B. and E.S. conceived the project and methodology. C.A.M.v.B and S.M.K. curated the data. K.O.O. implemented the model, performed the computational analysis and prepared the figures. E.Q., C.A.M.v.B, H.W.v.K, R.A.L.d.G., J.S.P.V., J.H.S.Y., M.A.N. performed wet lab experiments. E.S. supervised the study. K.O.O, C.A.M.v.B, and E.S. wrote the manuscript with feedback from H.V.

## Acknowledgments

We thank Leiden University Medical Center Flow Core Facility and Leiden Genome Technology Center for excellent purification of lymphoma cells and cell preparation and sequencing. We wish to express our deepest gratitude to all patients who allow us to use samples that are essential for our research.

## Supplementary Notes

### CaClust validation on simulated datasets

To test the performance of CaClust in a setting where the ground truth is known, and compare it with a simpler alternative model CACTUS [15], we devised simulation scenarios modelling three properties of an experimental FL dataset: the scRNA read depth, the number of BCR hyperclusters, and the variance of BCR sequences within a hypercluster. A scenario of median values was established as a baseline (referred to as *Basic*) and by varying one of these properties eight further scenarios were created with: i) high and ii) low scRNA reads; referred to as *High reads* and *Low reads*, respectively; iii) sparse clustering (resulting in more BCR hyperclusters, referred to as *Sparse clusters*); iv) high variance BCR sequences within hyperclusters (*High variance BCR*); and finally, four scenarios with centroid behaviour, where a portion *x* of cells within a fraction *y* of hyperclusters shares the most probable BCR sequence of its hypercluster’s BCR profile (in four versions of (*x, y*) v-viii): (0.8, 0.8), (0.8, 0.2), (0.2, 0.8), (0.2, 0.2); referred to as *x centroids in y clusters*). The simulation process and all scenario details are described in Methods. The data for each scenario was simulated 10 times.

CaClust outperforms CACTUS in all simulation scenarios, with a drastic increase in performance in some cases (Fig. 2). The major advantage of CaClust over CACTUS is particularly pronounced by the accuracy of assignment of cells to clones (Fig. 2a). In scenarios with at least basic scRNA read counts and basic or centroid BCR structure, CaClust achieves a near perfect accuracy. The concentration effect of CRP allows CaClust to reliably combine the scRNA information from cells with similar BCR sequences, resulting in a good accuracy even in the scenarios with low scRNA read counts. This effect is seen even in the scenarios with degraded BCR structure, i.e. sparse and high variance BCR hyperclusters. On the other hand, CACTUS performs well in scenarios with high number of scRNA reads but its performance quickly deteriorates in the basic and low reads scenarios, showing it does not utilise the information from hyperclusters with varied BCR sequences. It does however utilise the information from hyperclusters of cells with identical BCR, as shown by an improvement in performance in the simulation scenarios with centroid behaviour over the basic scenario.

In the task of genotype reconstruction CaClust shows a less pronounced improvement over CACTUS (Fig. **??**b). Both models perform near perfect in reconstructing the true clonal genotype profiles, with better reconstruction accuracy seen in scenarios with high scRNA read counts and worse accuracy in scenarios with degraded BCR structure.

For the hyperclustering reconstruction we see again major advantage of CaClust (Fig. **??**c). It achieves high adjusted rand index (ARI) scores in all but the high variance BCR scenario, which is specifically designed to give almost no BCR information. CACTUS performs poorly in all scenarios with basic and degraded BCR structure, since it does not manage to reconstruct hyperclusters with varied BCR sequences. However, in the scenarios with centroid behaviour, which should better represent the experimentaly observed BCR structure, it achieves at least a partial level of reconstruction.

### CaClust validation on patient data

We applied CaClust to real patient data, comprising of four patient samples: K4B, K5B, K6B and K7B. For these samples, we evaluated performance of CaClust to CACTUS by comparing their agreement with independent clustering of the cells by their gene expression profiles (see Methods) and by the uncertainty of assignment of cells to the clones.

To this end, three MCMC chains of both CaClust and CACTUS models were inferred on each sample until convergence (see Methods). After inference, we chose the CaClust and CACTUS chains with highest likelihoods for each sample.

Additionally, in the comparison with gene expression clustering we used two other approaches as a baseline: grouping cells with identical BCR sequences; and an ablation study with a stripped down version of the CaClust model, which only produces hyperclusters with no further grouping into clones. Importantly, these approaches are based purely on the BCR sequences and do not use scRNA variant data, so they can be treated as a more naive approach.

CaClust achieves better agreement with gene expression clustering as compared to CACTUS (Fig. 2d). This indicates that the clustering of cells into clones found using CaClust identifies clones that are more distinct on the phenotypic level. Moreover, both models achieve higher scores than the identical BCR grouping and the ablation study, showing that the integration of both BCR and scRNA variant data is more accurate in finding phenotypically distinct populations of cells.

Additionally, CaClust shows better certainty of cell assignment over CACTUS in all cases, demonstrating its better applicability on patient data (Fig. 2e).

### Model validation with targeted resequencing

As an additional validation of CaClust results, we performed an independent resequencing experiment, targeted for the variant sites in single cells (Methods). For samples K6B and K7B, 8 variant genes used in the model inference were included in single cell resequencing: *FCHSD2, TNFRSF14*, and *MFHAS1* from K6B; *AFF2, ANKRD12, CLSTN3, DNAJC11*, and *VMA21* from K7B. Their genotypes obtained from resequencing were compared with the the results of cell genotyping by CaClust.

However, in the resequencing results a high proportion of mutant cells showed monoallelic variant expression, as measured by the fraction of mutant cells with only mutant UMIs of a variant and no corresponding reference UMIs (pure mutant fraction, Supp. Tab. 2). Whether this is a result of the stochastic nature of gene transcription or the technical limitations of scRNA sequencing, in which not all transcripts are observed, it is a sign of a problem with cell genotyping based purely on the targeted resequencing. Specifically, for a given heterozygous variant, we expect the same number of cells with exclusively monoallelic alternative reads, as the number of cells with monoallelic reference reads for this variant, which would falsely classify them as wildtype. To account for this we proposed a simple correction for such random monoallelic expression (see next section and Supp. Fig. 2). In short, it modifies the total number of wildtype and mutant calls in resequencing to account for the possible misclassfied mutants. Importantly, the correction is agnostic to their CaClust genotypes to avoid bias (see next section). We applied the correction to heterozygous variants that showed any monoallelic variant expression in the resequencing data: *FCHSD2, TNFRSF14, CLSTN3, ANKRD12, DNAJC11*. Additionally, it must be noted that in the *VMA21* and *AFF2* homozygous mutations we also observed cells with a mixture of variant and mutated reads, which shows the inherent noise in sequencing experiments with amplification stages that could affect the agreement results.

We found that the VAF estimation from the corrected numbers of genotype calls in resequencing was in better agreement with the true VAF obtained from WES (Supp. Tab. 2). The VAF estimation from the resequencing results undervaluated the true VAF in all 8 variants, which hints at mutant cells being missed in initial resequencing calling. The correction application improved the VAF estimation for all variants, which stands as motivation behind its use, although in the case of a single gene *DNAJC11* it did result in an overvaluation of the VAF.

We calculated the agreement between resequencing and CaClust genotypes as the fraction of cells with the same genotype call on a variant position (mutated or wildtype). Despite the aforementioned inherent noisy nature of the scRNA sequencing data, CaClust achieved very high (*>* 90%) agreement for all 3 variants that did not qualify for the correction (homozygous *AFF2, VMA21*, or *MFHAS1* with no resequenced mutant reads). For the 5 heterozygous variants with possible missed mutant calls in the resequencing, we found 1 variant to be in high agreement before the correction, and 3 variants to be in high agreement after the correction (Supp. Tab. 2). Altogether, this shows the applicability of CaClust’s use for single-cell genotyping in FL and stands as additional validation for our model.

### Random monoallelic observation correction

In the analysis of the agreement between the model results and the resequencing genotype calling we apply a correction for the possibility of random monoallelic observations of the reference allele at heterozygous mutation positions, that accounts for dynamic autosomal random monoallelic expression (dynamic aRME) and the technical dropout of the sequencing experiments.

Maternal and paternal copies of genes are transcribed independently and in short intense bursts of transcription. Studies on allele-specific RNA transcription [50–53] have shown varying levels of RNA from maternal and paternal alleles within cell populations, with frequent observations of RNA from only one allele in a cell at a single point in time. In contrast to inherited aRME (clonal), this dynamic aRME is due to the stochastic nature of the gene expression process and is widespread across cell populations [50–52].

In our data we also observe monoallelic expression, both in the scRNA input data and the resequencing experiment, where a fraction of cells exhibits only variant reads of a heterozygous mutation, rather than a mix of variant and reference reads (with no CNA in that mutation’s region). A converse effect is therefore expected, in which only the reference allele was observed for a number of cells carrying the mutation. That poses a problem, since those cells would be misclassified in resequencing as non-carriers, thus inflating the False Positive (FP) and possibly True Negative (TN) numbers in mutation agreement between resequencing and CaClust.

As a solution we propose a simple correction for this random monoallelic observation (Supplementary Figure 4). Since the cells with a heterozygotic mutation harbour one copy of each of the alleles and each copy is expressed independently, we expect there to be a balance: if *M* cells are observed with purely mutated reads and *N* cells are observed with mixed variant and reference reads, we statistically expect further *M* mutated cells to have expressed only the reference allele and be misclassified by the resequencing genotype calling.

Of course, the distribution of those misclassifications between the FP and TN calls in agreement is not obvious; therefore, we distribute them proportionally.

The theoretical description of the correction is as follows. Let *M* be the number of cells in which only the mutated reads are observed and let *TP, FP, TN, FN* be the numbers of true positive, false positive, true negative and false negative initial calls in the agreement between resequencing and CaClust. Then we obtain the corrected agreement numbers *TP*^*′*^, *FP*^*′*^, *TN*^*′*^, *FN*^*′*^as described in Table 3.

**Table 3:** Application of a correction for random monoallelic expression with a balanced distribution.

| | if $M \leq (FP + TN)$ | if $M > (FP + TN)$ |
| --- | --- | --- |
| $TP'$ | $TP + \frac{FP}{FP+TN} \cdot M$ | $TP + FP$ |
| $FP'$ | $FP - \frac{FP}{FP+TN} \cdot M$ | 0 |
| $TN'$ | $TN - \frac{TN}{FP+TN} \cdot M$ | 0 |
| $FN'$ | $FN + \frac{TN}{FP+TN} \cdot M$ | $FN + TN$ |

## Supplementary Figures

**Supplementary Figure 1:**
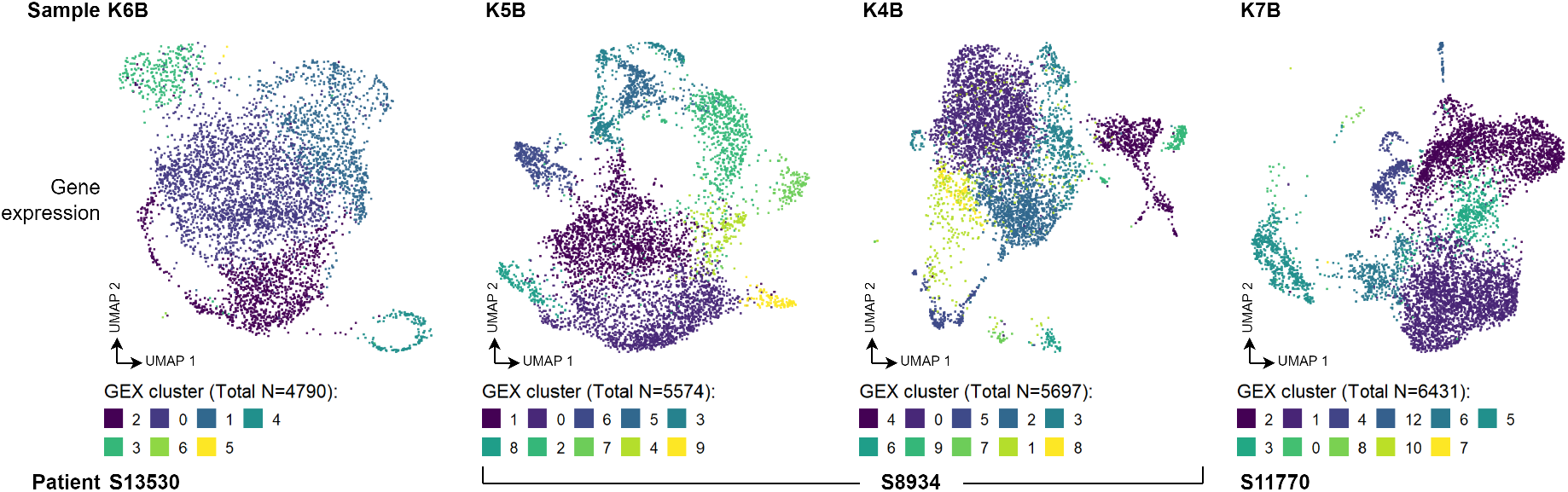
Clustering of cells by gene expression in each sample separately. Gene expression clusters were obtained by PCA dimensionality reduction to 30 dimensions and next Leiden clustering on the reduced profiles with resolution parameter of 0.3m using Seurat package.

**Supplementary Figure 2:**
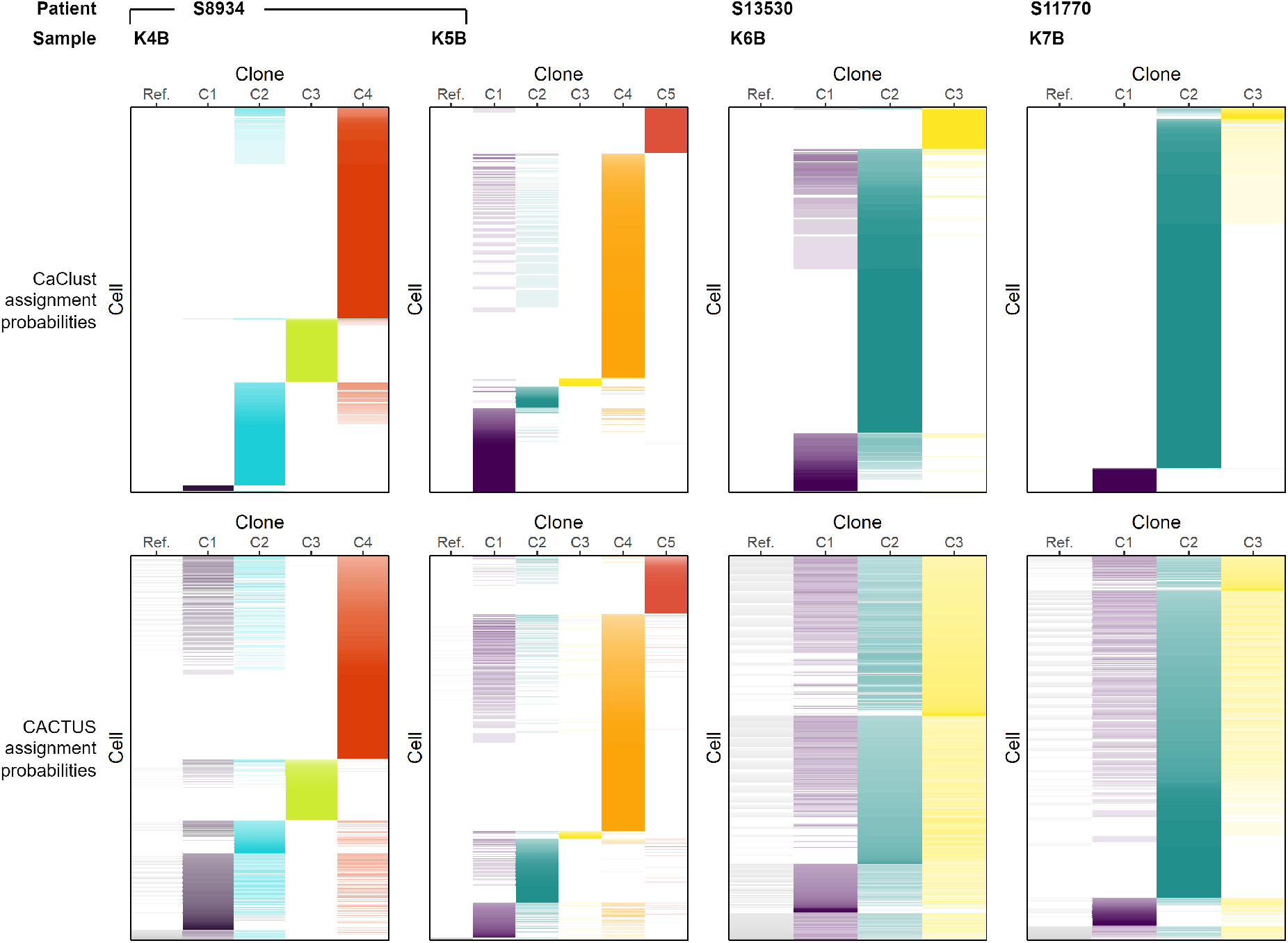
Heatmaps of assignment probabilities of cells to clones in all samples for CaClust and CACTUS. Cells order is not the same between methods. Clones are not shared between samples. Clone genotypes between methods are not identical for each sample; additionaly, CaClust clones C1 and C2 in each of the samples K4B and K5B are the counterparts of CACTUS clones C2 and C1 in those samples (resulting from the probabilistic nature of genotype correction in the methods).

**Supplementary Figure 3:**
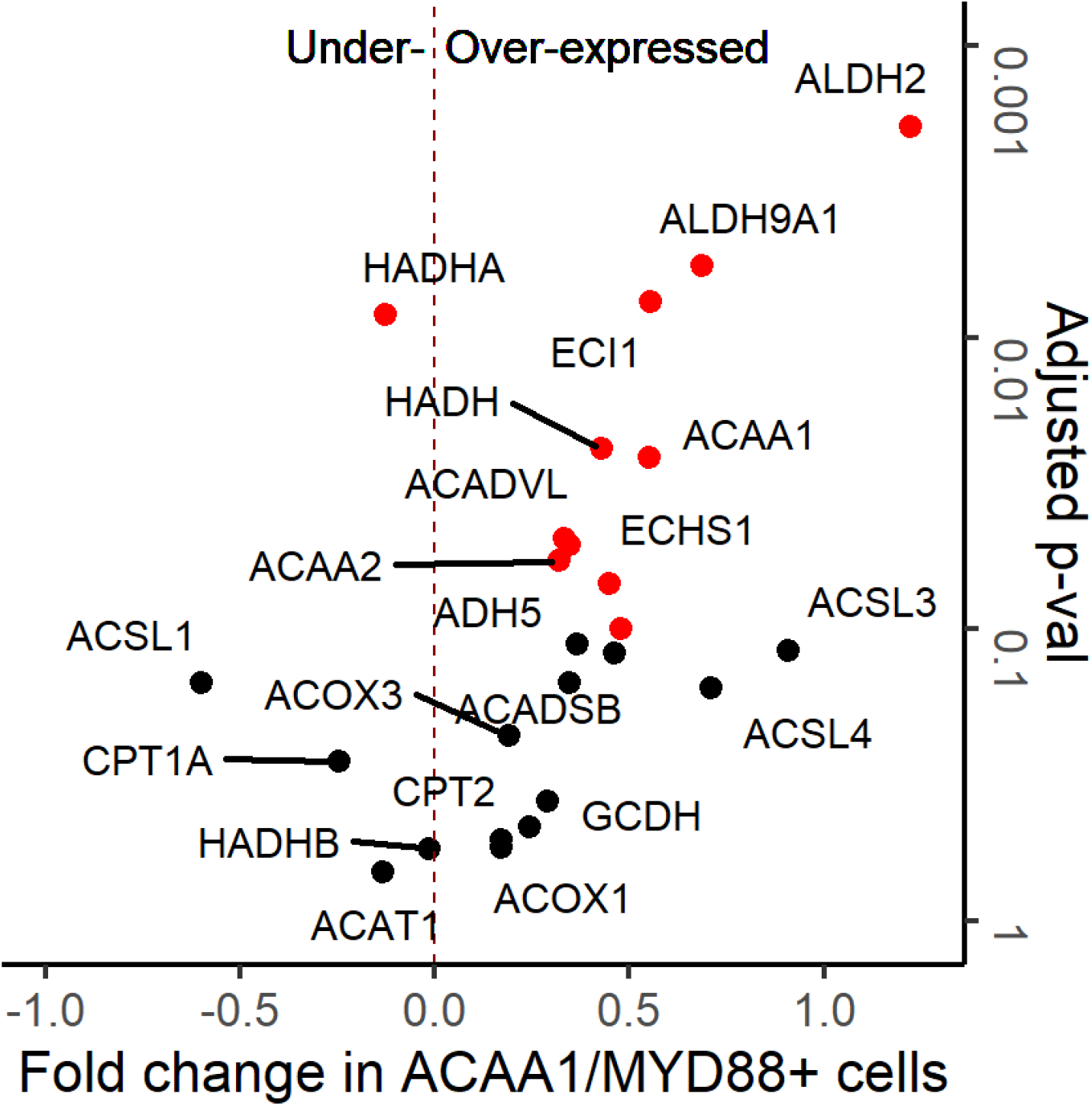
Enrichment of betaoxidation pathway genes in K7B clone C2. Genes with *p*_*adj*_ *<* 0.1 are highlighted.

**Supplementary Figure 4:**
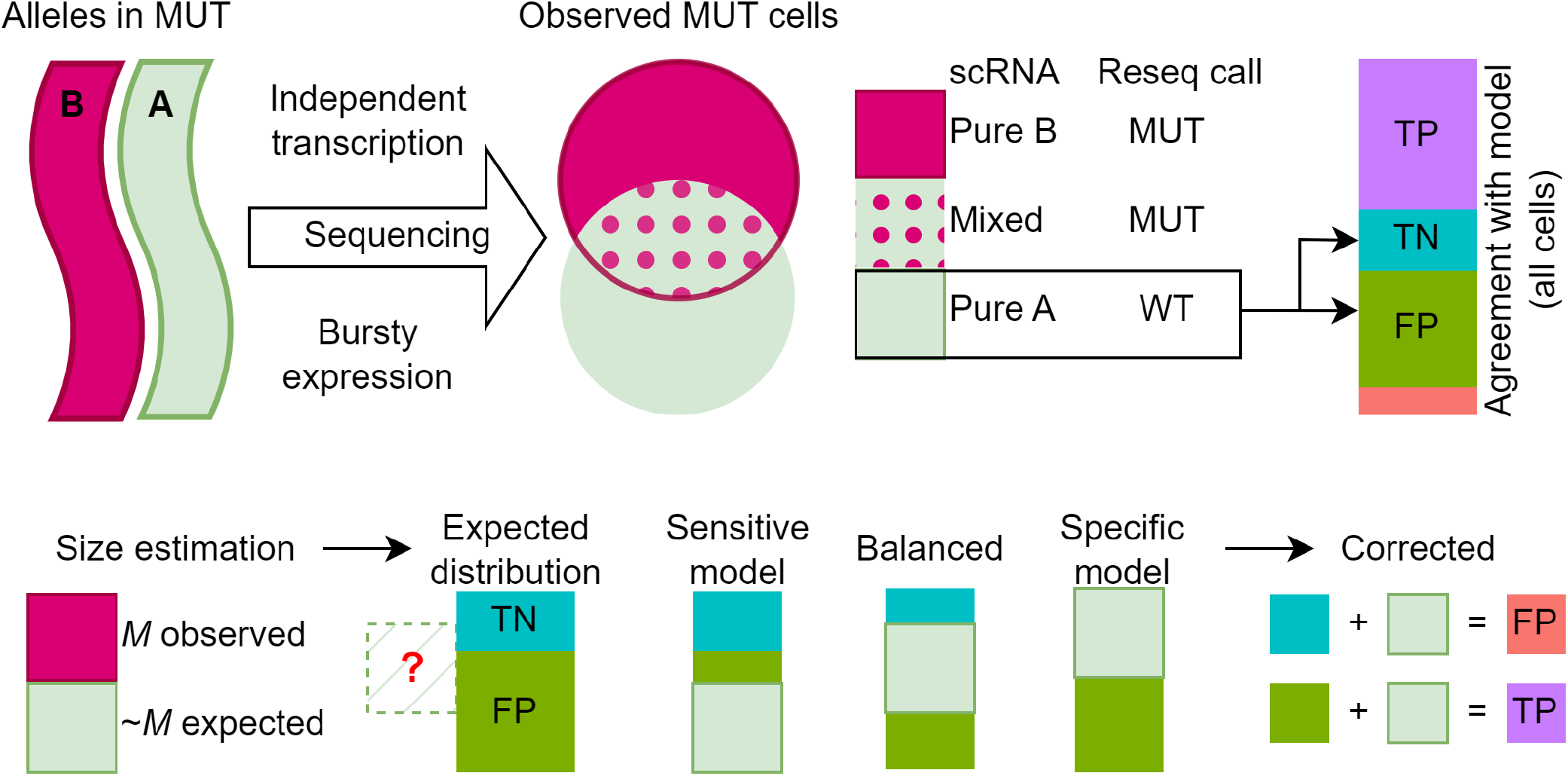
**Top**: illustration of the effect of stochastic expression of heterozy-gotic mutations on resequencing calling; heterozygous mutant cells with only reference scRNA reads in resequencing are falsely classified as wildtype, inflating the True Negative (TN) and False Positive (FP) agreement with calls from other methods. **Bottom**: Random monoallelic correction application. The effect size is estimated based on the number of cells with only mutant scRNA reads of the mutated position. The true distribution of the falsely classified cells between the TN and FP agreement calls is unknown and depends partially on the model’s sensitivity, to avoid bias we chose to apply a balanced correction in the paper.

